# A novel and critical role of the intracellular Zona Pellucida protein 2 (ZP2) for blastocyst formation in mice

**DOI:** 10.64898/2025.12.12.692802

**Authors:** Thomas Nolte, Steffen Israel, Hannes C.A. Drexler, Georg Fuellen, Michele Boiani

## Abstract

The zona pellucida (ZP) is the quintessential extracellular structure of mammalian oocytes. Contrary to long-standing view that the synthesis of ZP proteins is specific to oocytes and muted in embryos, we report here that the major zona pellucida protein ZP2 is re-synthesized and functionally required during mouse embryo development. The orthogonal methods of mass spectrometry and monoclonal immunofluorescence revealed an increase of ZP2 abundance at the 8-cell / morula stage, which did not occur when zygotes were microinjected with translation-blocking oligonucleotides (morpholinos). To shed light on the functional significance of embryonic ZP2, we performed protein knockdown using immunodepletion (by ‘Trim-Away’) while at the same time preventing replenishment (by translation-blocking morpholino). ZP2 knockdown resulted in morula stage retardation and formation of defective blastocysts, whose cell lineages trophectoderm and primitive endoderm were smaller and less able to support post-implantation development. The transcriptional correlates of these morphological alterations had a gene ontology (biological process) signature that included cell lineage-relevant terms (‘endoderm development’, ‘gastrulation’), while the proteomic correlates had a gene ontology signature related to protein synthesis. Taken together, these results call into question the traditional model that ZP proteins function solely in the extracellular space and accompany embryogenesis as passive bystanders: on the contrary, ZP proteins also participate actively in the intracellular processes of early embryogenesis.

## Introduction

The quintessential structure of mammalian oocytes, the zona pellucida, is known as the extracellular glycoproteic coat that envelopes the oocyte [1, 2] and that is synthesized only during oogenesis [3, 4]. In mice, which have three active zona pellucida genes (*Zp1, Zp2, Zp3*), the zona serves three known functions, namely, to encase the oocytes in the ovarian follicle, to mediate sperm-oocyte interaction while avoiding polyspermy, and to prevent premature contact between the embryo and the oviduct epithelium after fertilization. However, this model is likely incomplete. First, it does not explain why *Zp2* mutant (knockout, KO) mouse oocytes, with their abnormally thin zona that actually facilitates polyspermy, do not develop to term even after monospermic fertilization [5]. While the embryonic defect of *Zp2* KO oocytes [5] could be consequence of poor oocyte quality, developmental defects were also observed when zona proteins were disrupted in oocytes that had completed oogenesis normally and had been altered only thereafter [6–8]. Second, the model does not accommodate hints that the synthesis of ZP proteins may continue after oogenesis, as supported by the observation that *Zp* genes’ mRNAs are still found associated with polyribosomes of blastocyst-stage mouse embryos [9, 10]. Third, and consistent with the previous point, mass spectrometry studies revealed substantial amounts of ZP proteins present inside blastocysts, that is, intracellularly [11]. These amounts might be remnants of ZP proteins left unsecreted during oogenesis, or could hint at *de novo* protein synthesis during embryogenesis. We, therefore, evaluated the possibility that *de novo* protein translation of ZP2 takes place during embryogenesis to serve functions that are not encompassed in the current model.

Here we show that intracellular ZP2 is required for healthy blastocyst formation in mice. During zygotic cleavage, not only does ZP2 persist inside embryos throughout the pre-implantation phase, but its abundance also increases at the 8-cell / morula stage, as measured by mass spectrometry and direct immunofluorescence (IF) with monoclonal antibody. This increase is probably due to *de novo* ZP2 protein synthesis, since inhibition of *Zp2* mRNA translation with specific antisense oligonucleotides (morpholinos) prevents the increase. To illuminate the functional significance of the increase we performed protein knockdown (KD) of ZP2 using the method of immunodepletion by ‘Trim-away’ [12] reinforced by translation blockage (morpholino) [13] so as to prevent any replenishment of ZP2. ZP2 KD resulted in growth retardation at the morula stage, and resultant blastocysts presented a reduction in the size of the extraembryonic compartments (trophectoderm, primitive endoderm) as well as reduction in the rates of post-implantation development, compared to mock controls in which Trim-away and morpholino were directed to an irrelevant protein. Transcriptome and proteome analysis of the ZP2 KD blastocysts revealed alterations characterized by ‘endoderm development’ and ‘gastrulation’ as the gene ontology signature. These findings prompt new awareness about the biology of ZP proteins beyond their historical characterization as oocyte-specific gene products lacking active roles in embryogenesis.

## Materials and methods

### Ethical permits for animal experiments

Mice were used for experiments according to the license issued by the Landesamt für Natur, Umwelt und Verbraucherschutz of the State of North Rhine-Westphalia, Germany (license number LANUV 2024-320-Grundantrag), in accordance with the procedures laid down in the European Directive 2010/63/EU. We observed the ARRIVE guidelines [14] to the extent applicable.

### Mouse embryo production

All mice used in this study were reared in-house at the MPI Münster. They were maintained in individually ventilated type 2 L cages (EHRET GmbH Life Science Solutions, 79111 Freiburg, Germany), with autoclaved Aspen wood as bedding material and a cardboard tube as enrichment, in groups of five females or individually as males. Access to water (acidified to pH 2.5) and food (Teklad 2020SX, ENVIGO RMS GmbH, Düsseldorf, Germany) was *ad libitum*. The animal room was maintained at a controlled temperature of 22 °C, a relative humidity of 55 %, and a 14/10 h light/dark photoperiod (light on at 6:00 a.m.). The hygiene status was monitored every 3 months, as recommended by Federation of European Laboratory Animal Science Associations (FELASA), and the sentinel mice were found free of pathogens that might have affected the results of this study. Ovulation was induced by i.p. injection of pregnant mare’s serum gonadotrophin (eCG) (Pregmagon, IDT, 06861 Dessau-Roßlau, Germany) and hCG (Ovogest, MSD Tiergesundheit, Germany). Lean B6C3F1 females aged 8–10 weeks and weighing approx. 25 g were injected i.p. using a 27G needle, at 5 pm, with 10 I.U. eCG and 10 I.U. hCG 48 h apart. The females were mated to CD1 studs aged 3-12 months to produce zygotes, or left unmated to collect MII oocytes. On the next morning the females were killed by cervical dislocation. The cumulus-oocyte complexes were recovered, dissociated in hyaluronidase (cat. No. 151271, ICN Biomedicals, USA; 50 I.U./mL in HCZB), dissolved in Hepes-buffered Chatot, Ziomek and Bavister medium (HCZB) with bovine serum albumin (BSA; Probumin, Milllipore, Kankakee, IL, USA) replaced by polyvinylpyrrolidone (PVP, 40 kDa; cat. no. 529504, Calbiochem, EMD Biosciences, La Jolla, CA, USA) 0.1% w/v. The cumulus-free zygotes were cultured in 500 µL of Potassium (K) simplex optimization medium (KSOM) prepared in house as per original recipe [15]. KSOM contained free amino-acids (aa) both essential and non-essential (hence called KSOM(aa)), 0.2% (w/v) BSA and gentamicin (50 μg/mL), and was used in a four-well Nunc plate without oil overlay, at 37 °C under 6 % CO2 in air.

### Proteome analysis of all pre-implantation stages in data-dependent acquisition (DDA) mode

Embryos were collected for proteome analysis at consecutive stages up to blastocyst (n=100 embryos per sample). The zonae were either left in place or removed with acidic Tyrode’s solution. Following lysis of the embryos in PreOmics buffer (as part of the kit iST 8x; Preomics Planegg/Martinsried Germany), proteins were processed using the Preomics iST kit according to the manufacturer’s instructions (Preomics, 82152 Planegg-Martinsried, Germany). The digested and purified sample was subsequently dried in an Eppendorf Concentrator and resuspended in Buffer A (0.1% formic acid) for the liquid chromatography–mass spectrometry (LC-MS/MS) measuremen. Nanoflow reversed-phase liquid chromatography was performed using a nanoElute 2 UHPLC system (Bruker Daltonics), coupled online to a timsTOF Pro2 mass spectrometer via a CaptiveSpray nano-electrospray ion source (Bruker Daltonics). Peptide mixtures were chromatographically separated either on a 40cm long fused silica emitter (CoAnn Technologies) home-packed with 1.5 µm ReproSil Saphir 100 C18 beads (Dr. Maisch) or on a 50cm long PepSep column (Bruker)via a multisegment gradient from 5% Buffer B (80% acetonitrile, 0.1% formic acid) to 18% B within 57 min, from 18 – 27% B in 21 min, and from 27 – 37% B in 13 min, before being flushed with90% B (flow rate 300 nl/min). The column temperature was maintained at 50°C. The timsTOF was operated in the DDA mode using the DDA-PASEF standard_1.1_cycletime conditions (Bruker). Raw data were processed for identification and quantification by MaxQuant Software (version 2.0.3.0; [16]) with the ‘iBAQ’ option enabled and the ‘requantify’ option disabled. MaxQuant provides intensities for heavy and light labelled peptides and proteins separately and this also pertains to the iBAQ values. The search for identification was performed against the UniProt mouse database (version from 04/2019) concatenated with reversed sequence versions of all entries and supplemented with common laboratory contaminants. Parameters defined for the search were trypsin as the digesting enzyme, allowing two missed cleavages, a minimum length of six amino acids, carbamidomethylation at cysteine residues as a fixed modification, oxidation at methionine, and protein N-terminal acetylation as variable modifications. The maximum mass deviation allowed was 20 ppm for the MS and 0.5 Da for the MS/MS scans. Protein groups were regarded as identified with a false discovery rate (FDR) set to 1 % for all peptide and protein identifications; only proteins with at least 2 identified unique peptides were considered further. The molar fractional content of each protein P in a sample (relative iBAQ intensities = riBAQP) was determined for the light and heavy labeled proteoforms independently, according to Shin *et al.* (2013), as follows:

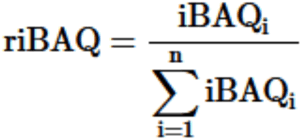

The mass spectrometry proteomics data have been deposited in the ProteomeXchange Consortium via the PRIDE repository (http://proteomecentral.proteomexchange.org), with the dataset identifier PXD056725.

### Evacuation of zonae for immunoblot analysis

Oocytes were transferred in groups of 10 to a micromanipulation drop on the stage of a Nikon Eclipse TE2000-U inverted microscope fitted with Nomarski optics and Narishige micromanipulator. The micromanipulation medium consisted of HCZB. Each oocyte was held firmly in place with the holding needle by applying negative pressure (suction). An Eppendorf’s TransferTip (ES) needle operated by an Eppendorf’s CellTram Vario (Eppendorf, Hamburg, Germany) was used to pierce through the zona and aspirate the oocyte. The evacuated zonae were concentrated on the bottom of a tube by centrifugation, the supernatant was replaced with RIPA buffer, and the lysate was processed further as described below.

### Immunofluorescence (IF) analysis of ZP2 in oocytes and pre-implantation embryos

Oocytes or embryos were analyzed by performing an immunostaining followed by confocal microscopy imaging, as per our routine protocol. Briefly, embryos were fixed with 1.5 % formaldehyde in 1x PBS for 20 min at 4 °C, permeabilized with 0.1% Triton X-100 in 1x PBS for 20 min at 4 °C, and then blocked with 2 % BSA, 2 % glycine, 5 % donkey serum, 0.1 % Tween20 in 1x PBS overnight at 4 °C. When applicable, the zona was removed by bathing the oocytes or embryos in acidic Tyrode’s solution (cat.no. T1788, Sigma-Aldrich Chemie GmbH, 82024 Taufkirchen Germany) prewarmed at 30 °C. The primary anti-ZP2 antibody was monoclonal IgG raised in rat-mouse hybridoma (CRL-2463, ATCC IE-3) applied at 1:200 overnight at 4 °C. An appropriate Alexa Fluor-tagged secondary antibody (Invitrogen, Thermo Fisher Scientific, Karlsruhe, Germany) was matched to the primary and incubated for 1-2 h at room temperature. Direct labeling of IE antibody: Hybridoma anti-ZP2 antibody ATCC IE-3 CRL-2463 was used for direct labeling. The labeling was performed after the instruction of the Alexa Fluor® 647 Conjugation Kit (Fast)- Lightning-Link® (Abcam cat.no. ab269823). DNA counterstaining was performed with YO-PRO-1 or Hoechst 33342 (1 µM). For imaging, embryos were placed in 5 µl drops of PBS on a 50-mm thin-bottom plastic dish (Lumox hydrophilic dish, Greiner Bio-One, Frickenhausen, Germany) and overlaid with mineral oil (cat. no. M8410, Sigma-Aldrich Chemie GmbH, 82024 Taufkirchen, Germany). Images were captured using a 20x CFI Plan Apochromat VC objective on an inverted motorized Nikon TiE2000 microscope fitted with an Andor Dragonfly 502 spinning disc confocal unit Scanning System. Optical sections per embryo were captured using a high resolution (2048 x 2048 pixel) sCMOScamera. Maximum projections were analyzed with Fiji [17].

### High-resolution immunofluorescence imaging of ZP2 subcellular localization with Airyscan

The analysis of ZP2 distribution and localization in pre-implantative mouse embryos were performed by direct and indirect IF. Therefore, embryos were fixed with 1.5 % formaldehyde in 1x PBS for at least 30 min at 4 °C. The embryos were then permeabilized with 0.1 % Triton X-100 in 1x PBS for 20 min at 4 °C and blocked with 2 % BSA, 2 % glycine, 5 % donkey serum and 0.1 % Tween20 in 1x PBS overnight at 4 °C. After blocking the zona was removed by Tyrode solution. For the direct and indirect IF, the primary ZP2 antibody ATCC IE3 CRL-2463 was applied at 1:200 and incubated overnight at 4 °C. Hoechst 33342 was used for DNA counterstaining. Super-resolution images were captured by using a 20x Plan-Apochromat objective on a laser scanning confocal microscope (LSM980) with the Zeiss Zen (Zeiss Zen version 3.5) software. Images were acquired using Airyscan 2 detector.

### Immunoblotting analysis of ZP2 expression in pre-implantation mouse embryos

Oocytes or embryos were gently centrifuged in protein-free HCZB medium at 100 G for 20 min to form a tiny pellet. The supernatant was carefully aspirated using a mouth-operated micropipette, and replaced by RIPA buffer containing protease inhibitors. The resultant lysates were mixed with 6x Laemmli sample buffer and boiled for 5 min at 99 °C. These samples were loaded on a 12 % separation gel and blotted onto a PVDF membrane. The membrane was blocked for at least 3 h and incubated (3 % nonfat dry milk in 0.1 % PBS-Tween20) with primary antibodies overnight at 4 °C. The antibodies against ZP2 (cat.no. A10126, Abclonal; IE-3, ATCC cat.no. IE-3 – CRL-2463; 21832-1-AP, ProteinTech) were applied at a dilution factor of 1: 2000. Signal intensities were standardized on α-tubulin (1:5000, cat. no. T6199, Sigma-Aldrich Chemie GmbH). After 3X washing in 0.1 % PBS-Tween20, the blot was incubated with horse-radish peroxidase (HRP)-coupled secondary antibody at room temperature for 1 h. The membrane was washed and then developed with chemiluminescent HRP substrate solution. The chemiluminescent signal was detected using the AGFA Curix 60.

### Synthesis and purification of *mCherry-Trim21* mRNA

mRNA was synthesized from an expression construct (pGEMHE-*mCherry*-*trim21* plasmid, Addgene Plasmid #105522). The plasmid was linearized with SwaI (cat. no. FD1244, Thermo Fisher Scientific, Karlsruhe, Germany). Capped mRNA was synthesized with T7 polymerase (Ambion mMessage mMachine T7 kit) according to manufacturer’s instructions. The *mCherry-Trim21* mRNA was purified with Quick-RNA MicroPrep (cat. no. R1051, Zymo Research, 79110 Freiburg, Germany) and preserved in MilliQ water at -80 °C.

### Synthesis and purification of ZP2 antibody

The antibody was produced in-house from hybridoma cells (ATCC collection). The antibody was washed three times and concentrated at 4 °C using Amicon Ultra-0.5 100 kDa centrifugal filter devices (cat. no. UFC100, Merck Millipore, Merck Millipore, Darmstadt, Germany), which remove salts and impurities, allowing to elute the antibody in pure water.

### Synthesis of morpholino oligonucleotides

Translation-blocking morpholino oligonucleotides, 25 nucleotides in length, were purchased from Gene Tools, LLC (Philomath, OR 97370, USA). They were dissolved in MilliQ water at 1 mM and aliquoted before storing frozen. Sequences were selected by design parameters according to the manufacturer’s recommendations, namely approximately 50% G/C and A/T content with no predicted hairpins. Morpholinos were designed for blocking translational initiation by binding to sequences flanking the AUG initiating methionine of the two isoforms of the *Zp2* gene (Ensemble accession number ENSMUST00000033207.6 (Zp2-201) and ENSMUST00000208874.2 (Zp2-205)). The following oligonucleotide sequences were used. Morpholino targeting ZP2: 5‘ – CTCTGCCACCTCGC<u>CAT</u>GTTGGAAG– 3‘, suited to bind both ZP2 isoforms and almost identical to the oligo previously used to inhibit ZP2 protein synthesis during oogenesis (5‘ – CCACCTCGC<u>CAT</u>GTTGGAAGGTAC – 3‘; [18]); morpholino targeting GFP: 5‘ – ACAGCTCCTCGCCCTTGCTCACCAT– 3’. The CAT sequence complementary to the start codon is underlined in the case of a molecule binding to a sequence containing the AUG. Using homology searches (BLAST search), we also systematically tested each sequence for possible interaction with other sequences within the mouse genome, finding none except the *Zp2* gene.

### Knockdown (KD) of ZP2 using *Trim-Away* combined with morpholino

Zygotes were microinjected with a cocktail of *mCherry-Trim21* mRNA, Oregon Green dextran beads (OGDB) 70 kDa, anti-ZP2 antibody (ATCC cat.no. IE-3 – CRL-2463) and *Zp2* morpholino at approximately 16 h post-hCG. Concentrations in the cocktail were: 0.15 mg/mL mRNA, 0.017 mg/mL OGDB, 5 mg/mL antibody, and 0.01 mM morpholino, respectively, in MilliQ water. As a mock KD control, ZP2 antibody was replaced with GFP antibody (Thermo Fisher G10362) and *ZP2* morpholino was replaced with anti-GFP morpholino, since GFP is not expressed in the wild-type embryos used in this study. Microinjection of the mRNA-antibody-OGDB mixture was conducted on the stage of a Nikon TE2000U microscope fitted with a piezo drill (PrimeTech, 635 Nakamukaihara Tsuchiura Ibaraki 300-0841 Japan), using a blunt-end glass needle (inner diameter 6-7 microns, outer diameter 8-9 microns) filled with 2-3 µl mercury at the tip. Volumes were pressure-injected into the zygote using a Gilmont GS-1200 micrometer syringe operated manually. During the microinjection, cells were kept in a 200-300 µl drop of HCZB medium on a glass-bottomed (Nomarski optics) dish at a room temperature of 28 °C. After microinjection, zygotes or embryos were allowed to recover in the drop for 5–10 min, before returning them to KSOM(aa) medium.

### Analysis of cell lineage allocation of blastocysts

Blastocysts were fixed as described (“Immunofluorescence (IF) analysis of ZP2 in oocytes and pre-implantation embryos”) and subjected to triple immunostaining followed by confocal microscopy imaging to identify and map the different cell lineages, as described previously [19]. The following primary antibodies were applied simultaneously to the specimens overnight at 4 °C for analysis of cell lineage allocation: anti-CDX2 mouse IgG1 (Emergo Europe, The Hague, Netherlands, cat. no. CDX2-88), anti-NANOG rabbit IgG (Cosmo Bio, Tokyo, Japan, cat. no. REC-RCAB0002P-F) and anti-SOX17 goat IgG (R&D Systems, cat no. AF1924) in dilutions of 1:200, 1:2,000 and 1:100, respectively. Appropriate Alexa Fluor-tagged secondary antibodies (Invitrogen) were matched to the primaries and incubated for 1–2 h at room temperature. Blastocysts were imaged as described (“Immunofluorescence (IF) analysis of ZP2 in oocytes and pre-implantation embryos”). Cells were counted in maximum projection images using Image-J.

### Outgrowth assay

We transferred blastocysts individually onto a feeder layer of γ-ray-inactivated (30 *Gray)* mouse embryonic fibroblasts (C3H background) grown to confluence in 96-well plates (flat bottom) previously, using our adaptation [20] of the outgrowth method [21]. The medium consisted of high-glucose Dulbecco’s modified eagle *medium* with high glucose (CAT No. D5671, Sigma-Aldrich Chemie GmbH) supplemented with 15 % heat-inactivated fetal bovine serum (BioWest, Nuaillé, France), GlutaMAX 1X (CAT. No. 35050-038, Gibco at Thermo Scientific), penicillin/streptomycin 1X (CAT. No. P4333, Sigma-Aldrich Chemie GmbH), nonessential amino acids 1X (CAT. No. M7145, Sigma-Aldrich Chemie GmbH), mercaptoethanol 0.1 mM (CAT No. 31350-010, Gibco at Thermo Scientific) and 1000 Units/mL LIF (produced in-house).

### Transfer of mouse embryos to the pseudopregnant females

Embryos were surgically transferred in groups of ten to the genital tract of pseudo-pregnant CD1 recipients. We used two schemes. In one scheme, we transferred 4-cell embryos into one oviduct of females had received the copulation plug from vasectomized CD1 males on the same day as the embryo transfer. In the other scheme, we transferred blastocyst-stage embryos into both uterine horns of females had received the copulation plug from vasectomized CD1 males two days prior to embryo transfer. Prior to surgery, CD1 foster mothers were anesthetized with Ketamine (80 mg/kg body weight)/Xylazin (16 mg/kg)/Tramadol (15 mg/kg) in PBS, delivered i.p. The surgical wounds were sutured with resorbable Marlin violett. Post-surgical pain was alleviated by providing the animals with Tramadol in drinking water (1 mg/mL). Pregnancies were evaluated by C-section shortly before term (embryonic day 16.5).

### Transcriptome analysis of blastocyst-stage embryos after ZP2 KD

Sequencing and raw data processing was provided by the Core Genomic Facility of the Faculty of Medicine of the University of Muenster, as follows. RNA was extracted and purified using Quick-RNA MicroPrep (cat. no. R1051, Zymo Research Europe GmbH). The library preparation of the total RNA was performed with the Watchmaker mRNA Library Prep Kit (7K0078-024) according to the manufacturer’s instructions. Paired-end sequencing with a read length of 111bp was performed on NovaSeq® X Plus System using the corresponding NovaSeq® X Series 1.5B Reagent Kit (200 Cyc). Total RNA integrity and quality of the library were assessed using a TapeStation4200 (Agilent, Santa Clara, CA, USA). On average, the libraries contained 24.7 ± 2.9 million 111-base paired-end reads. Using a molecular barcode, the samples were demultiplexed (DRAGEN BCL Convert 4.3.13) to fastq data and quality controlled (FastQC version 0.12.1; https://www.bioinformatics.babraham.ac.uk/projects/fastqc/). Trimmomatic (PMID 24695404; version 0.39) was used for adapter trimming and read filtering. Reads were aligned to the reference genome (Mus musculus Ensembl GRCm39) using Hisat2 (PMID 25751142; version: 2.2.1). Aligned reads were sorted using samtools (PMID 21903627; version 1.16.1). Sorted and aligned reads were counted into genes using HTSeq framework (PMID 25260700; version 2.0.3), then these counts were used to calculate the FPKM to normalize for sequencing depth and gene length. StringTie version 3.0.0. was used to calculate the potential transcripts. FPKM values for Ensembl gene identifiers corresponding to the same gene symbol were averaged, and the values associated with each gene symbol were averaged across replicates. The RNA-seq data have been deposited in the ArrayExpress repository, with the dataset identifier E-MTAB-16339.

### Proteome analysis analysis of late morula-stage embryos in data-independent (DIA) acquisition mode after ZP2 KD

Morulae were collected from each group on day 3, deprived of the zona pellucida by washing in acidic Tyrode solution, lysed in PreOmics iST LYSE buffer and stored at -20°C until further processing. Lysates were processed using the PreOmics iST kit according to the manufacturer’s protocol. Purified peptides were dried down in an Eppendorf Concentrator, eventually dissolved in 10 µl LC-LOAD loading buffer, of which 1 µl was injected into the mass spectrometer. Nanoflow reversed-phase liquid chromatography was performed using a nanoElute 2 UHPLC system (Bruker Daltonics), coupled online to a timsTOF Pro2 mass spectrometer via a CaptiveSpray nano-electrospray ion source (Bruker Daltonics). Peptide separation was achieved using a 25 cm × 75 µm fused silica capillary column (CoAnn Technologies), home-packed with 1.5 µm ReproSil Saphir 100 C18 beads (Dr. Maisch). To elute bound peptides a linear gradient of 4–27% buffer B (99.9% acetonitrile, 0.1% formic acid) over 60 minutes was applied, with buffer A consisting of 0.1% formic acid. The flow rate was set to 300 nl/min. The mass spectrometer was operated in DIA-PASEF mode using the standard DIA-PASEF long-gradient method. Signals were recorded across a mass range of 100–1700 m/z and an ion mobility range from 1/K₀ = 0.60 to 1.60 Vs/cm², with a ramp time of 100 ms in the dual TIMS analyzer. Collision energy was adjusted as a function of ion mobility, ranging from 59 eV at 1/K₀ = 1.60 Vs/cm² to 20 eV at 1/K₀ = 0.6 Vs/cm².

Raw mass spectrometry data were processed using DIA-NN software (v2.0.2) in library-free search mode with heuristic protein inference performed at the gene level. Neural networks were employed in single-pass mode, enabling matching between runs, and QuantUMS was applied as the quantification strategy. The predicted library included entries sourced from mouse UniProt UP000000589_10090_additional.fasta database (April 2024 version), including isoforms, the mCherry sequence, and common laboratory contaminants. Search parameters included carbamidomethylation of cysteine residues as a fixed modification, while oxidation of methionine and acetylation of protein N-termini were set as variable modifications. Trypsin was specified as the digesting enzyme, allowing up to two missed cleavages and peptides with lengths between 7–30 amino acids. Mass tolerances were set to 20 ppm for MS scans and 15 ppm for MS/MS scans, with charges above +4 excluded from analysis. Protein groups were considered identified with a false discovery rate (FDR) of ≤1% at both peptide and protein levels. Initial downstream analysis was performed using Perseus software (v1.6.15.0). Common laboratory contaminants were removed from the dataset, and protein intensity values provided in the pg_matrix.tsv table were log₂-transformed. The mass spectrometry proteomics data have been deposited in the ProteomeXchange Consortium via the PRIDE repository (http://proteomecentral.proteomexchange.org), with the dataset identifier PXD062983.

### Principal component analysis (PCA) and over-representation analysis of differentially expressed mRNAs and proteins

RNA sequencing (RNA-seq) data analysis was performed in-house using the output of the NovaSeq® X Plus System, exported to Microsoft Excel (Microsoft Deutschland GmbH, 80807 München, Germany) format and imported in JMP Pro v. 18 (JMP Austria, Germany & Switzerland, 69188 Heidelberg, Germany). Proteome data analysis was performed in-house using the output of the MaxQuant or Perseus software exported to Microsoft Excel. For the PCA, we considered the transcripts or proteins that were present in all samples of all replicates. The gene lists were uploaded to JMP v.18 to generate the PCA. Differentially expressed transcripts or proteins were identified in Volcano plots using p < 0.05 (Student’s t-test) and a |fold change|>2. The number of transcripts or proteins was further narrowed down using Venn diagrams to isolate genes or proteins that are differentially expressed exclusively in ZP2 KD or mock KD but not in the intersection. Over-representation analysis was performed with the WEB-based GEne SeT AnaLysis Toolkit (*Webgestalt*) [22, 23]. We selected the ontology ‘Biological process’ (BP). Terms with a FDR ≤ 0.05 were considered enriched.

### Statistical analysis of developmental rates and IF images

Developmental rates of embryos and image intensities of IF were analyzed non-parametrically by Wilcoxon test using the statistical program JMP v.18 (JMP Austria, Germany & Switzerland, 69188 Heidelberg, Germany). Transcript and protein levels of ZP2 KD, mock KD and micromanipulation control were compared with Student’s t-test using Microsoft Excel.

## Results

### Unexpected increase in intracellular ZP proteins abundance at the 8-cell / morula stage of mouse embryo development

We sought to understand whether ZP2 proteins decrease monotonically from oocyte to blastocyst, as would be expected of maternal gene products that are active only during oogenesis [3, 24], or whether they increase, as suggested by ribosequencing studies [9, 10], which measure the binding of transcripts to ribosomes and therefore their availability for translation. To this end we examined naturally fertilized mouse embryos using two orthogonal methods for measuring protein, namely, mass spectrometry (MS) and immunofluorescence (IF).

Using MS, we interrogated the proteomes of two developmental series (oocyte, zygote, 2-cell, 4-cell, 8-cell, morula, blastocyst). In one series the zona was left intact, in the other series it was removed with acidic Tyrode’s solution immediately prior to sample preparation for MS. This removal facilitates the detection of variations in intracellular protein content, since the extracellular zona accounts for 17% of the total protein content of a mouse oocyte [25]. Proteins were quantified in data-dependent acquisition mode using the ‘intensity-based absolute quantification’ (iBAQ) algorithm [26, 27]. iBAQ values are approximately proportional to the number of moles of protein present and thus iBAQi/ΣjiBAQj (adimensional) is the relative molar amount of protein ‘i’ among all proteins ‘j’, called relative iBAQ, briefly riBAQ [28]. The data of both series were concordant in showing that the ZP proteins undergo a transient increase at the 8-cell stage (Figure 1A). On the contrary, this profile is not seen in the projections of housekeeping genes (*Gapdh*, *Pgk1*, *Tubulin*, *Hprt*) (Figure 1B), suggesting that the variation in ZP proteins is specific.

**Figure 1.**
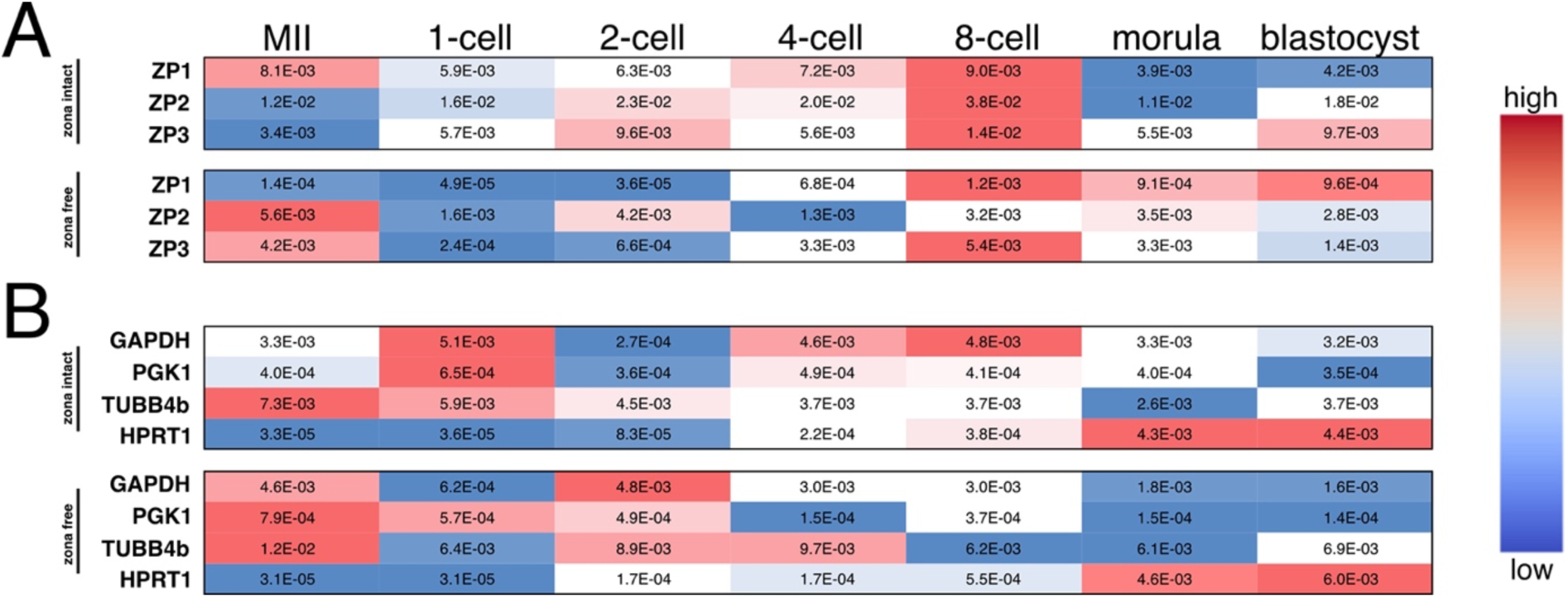
Mass spectrometric (LC-MS/MS) quantification of ZP protein levels during mouse pre-implantation development. Mouse embryos were collected at consecutive stages of pre-implantation development *in vitro* after culture in KSOM(aa) medium, and subjected to LC-MS/MS. Zona pellucida was either left in place (zona intact) or removed (zona free). Shown are the relative iBAQ (riBAQ) values of protein abundance for (A) ZP proteins and (B) housekeeping proteins. The color coding, indicative of expression level, was applied in each row independently of the other rows. The raw proteomic data are available in PRIDE repository with accession number PXD056725 and are provided in simplified form in Supplementary Table S1. Abbreviations: riBAQ, relative intensity Based Absolute Quantification [26, 27]; LC-MS/MS, Liquid Chromatography with tandem mass spectrometry. Abbreviations: iBAQ, intensity-based absolute quantification; LC-MS/MS, Liquid Chromatography with tandem mass spectrometry; MII, metaphase II oocyte.

Using the orthogonal method of IF and focusing on ZP2, we examined three developmental series of zygotes. One series was cultured *in vitro* and subjected to indirect IF (primary antibody + labeled secondary antibody). A second series was cultured *in vitro* and subjected to direct IF (labeled primary antibody). Since culture conditions can induce gene expression artefacts [29], a third series was obtained fresh from the genital tract, without *in vitro* culture, and was examined by indirect IF. In each series the different stages were immunostained simultaneously. A suitable antibody was identified by comparing three commercially available antibodies directed against different regions of ZP2 protein (ATCC cat.no. IE-3; ABclonal cat.no. A10126; ProteinTech cat.no. 21832-1-AP; Supplementary Figure S1A). Only IE-3 produced a single prominent band of the expected molecular weight, and this band was stronger in lysates of zona-intact oocytes compared to lysates of evacuated zonae and zona-free oocytes (with no signal in lysates of MEFs and ES cells; Supplementary Figure S1B). We also verified that IE-3 produced the same result, except of course in terms of intensity, after direct or indirect IF (Supplementary Figure S2). In all three series the ZP2 signal intensity declined after fertilization, increased at the 8-cell / morula stage, and then declined again at the blastocyst stage (Figure 2).

**Figure 2.**
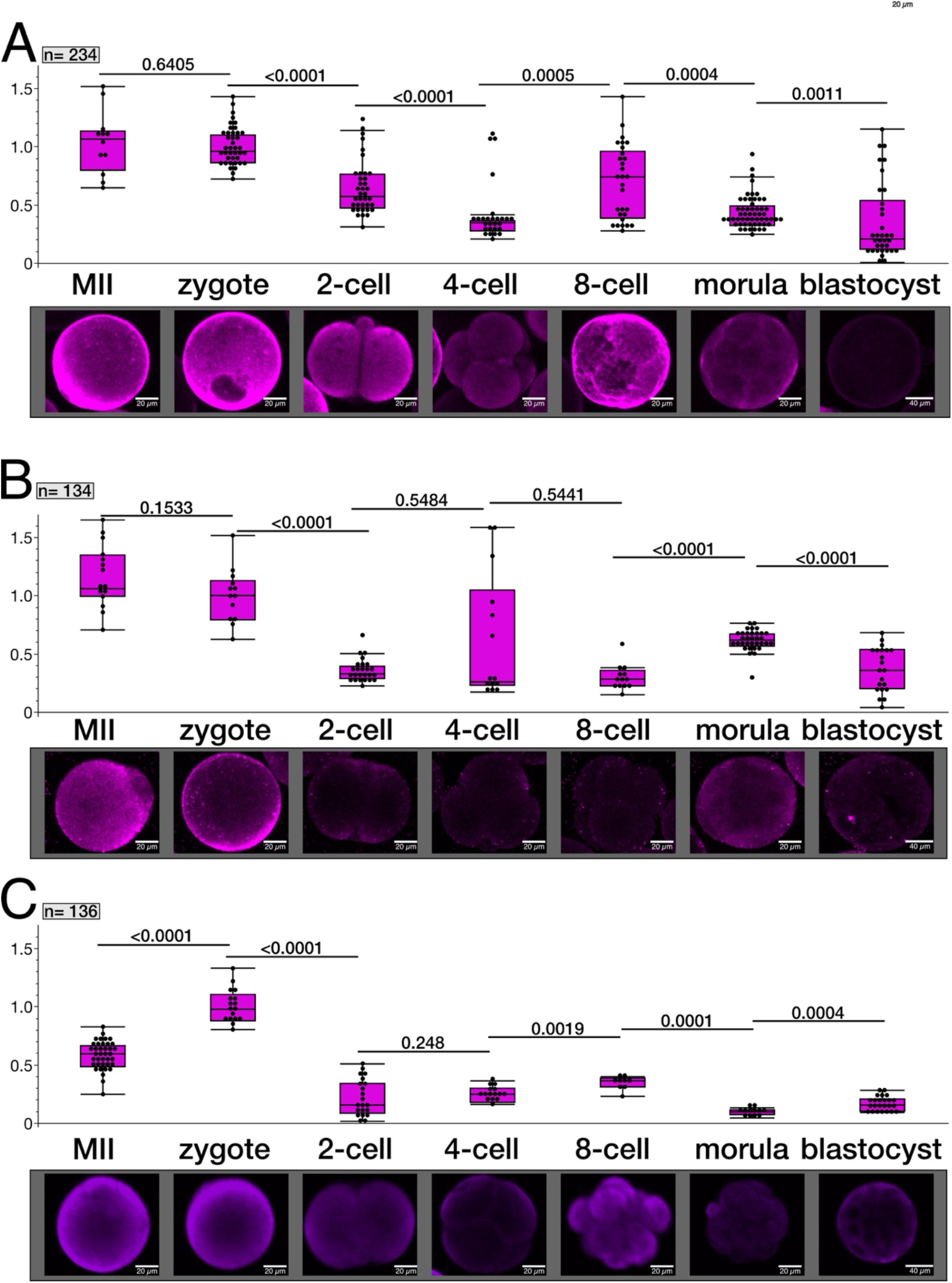
Immunofluorescence profiling of ZP2 protein during mouse pre-implantation development. (A) Developmental series of embryos cultured *in vitro* and subjected to indirect IF. (B) Developmental series of embryos cultured *in vitro* and subjected to direct IF. (C) Developmental series of embryos obtained fresh from the genital tract without *in vitro* culture and examined by indirect IF. Numbers shown in the top left corner of each diagram indicate the total numbers of embryos examined across all stages. The integrated density values of the ZP2 IF intensities were quantified using Image-J, normalized to the 1-cell stage (set to 1) and plotted by stage of development. Each point corresponds to the normalized integrated density one embryo. Each point corresponds to the normalized integrated density one embryo. All normalized integrated density values are provided in Supplementary Table S2. P-values were calculated with Wilcoxon test. Abbreviations: IF, immunofluorescence; MII, metaphase II (oocyte).

The results of these experiments conducted with orthogonal methods converge on an *interim* conclusion: the ZP2 protein, considered extracellular and oocyte-specific, is instead present inside embryos, and moreover, it increases at mid-preimplantation. Unclear is how the observed increase of ZP2 during embryogenesis comes about.

### Intracellular increase of ZP2 protein is contributed by *de novo* protein synthesis

Given the ZP2 protein increase observed at the 8-cell / morula stage, we sought to determine the origin of the increase. This may come from the transcription of new mRNAs or from the translation of pre-existing mRNAs. However, reanalysis of our previous RNA-seq dataset [11] showed that *Zp2* mRNAs decline monotonically from zygote to blastocysts (Supplementary Figure S3), whereas *Zp2* transcripts are known to be bound to ribosomes and thereby available for translation [9, 10]. We reasoned that if translation were involved, then blocking the translation of *Zp*2 mRNA should prevent the increase observed at the 8-cell / morula stage (Figure 1, Figure 2). Zygotes were thus microinjected with a morpholino oligonucleotide that spans the translaäon iniäaäon codon of *Zp2* mRNA and was previously used to inhibit ZP2 translaäon during oogenesis [18]. As a negaäve control, zygotes were microinjected with a morpholino targeting an irrelevant sequence, such as that of the green fluorescent protein (GFP), which is not present in the wildtype embryos of this study. A titration was performed to identify the optimal morpholino dose that would not compromise blastocyst formation of microinjected zygotes (Figure 3A). Using this dose (0.01 mM), the resulting 8-cell embryos presented a ∼ 20% reduction of ZP2 immunofluorescence intensity after they received the ZP2 morpholino compared to the negative control (Figure 3B). The results of these morpholino experiments support that the intracellular increase of ZP2 protein detected at the 8-cell / morula stage is contributed by *de novo* protein synthesis. It remains unclear however whether the intracellular ZP2 is a spurious occurrence or serves a functional purpose.

**Figure 3.**
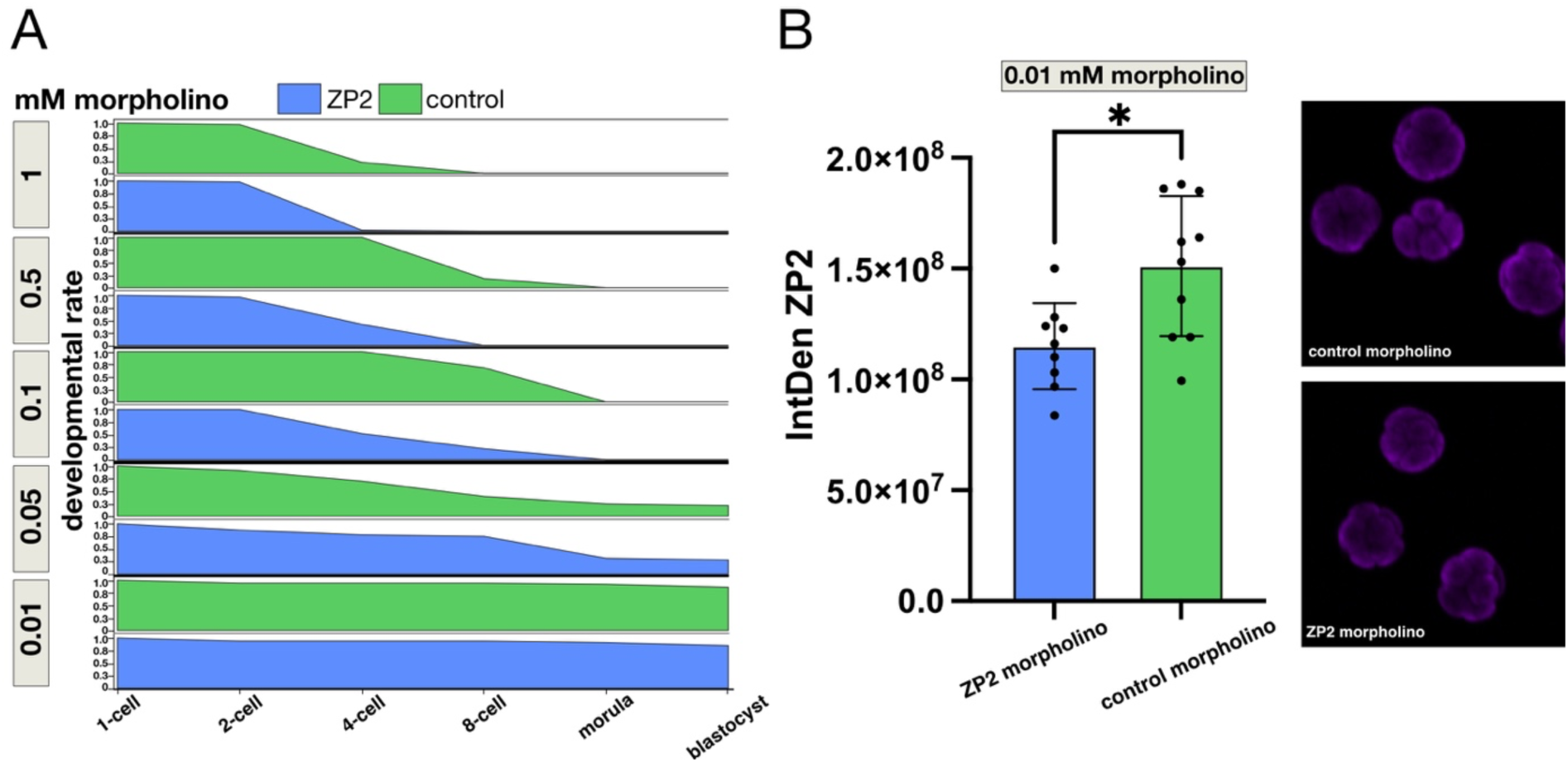
Intracellular increase of ZP2 protein abundance is contributed by *de novo* protein synthesis. (A). Developmental progression of zygotes microinjected with various concentrations of morpholino targeting *Zp2* mRNA (treatment) or morpholino targeting an irrelevant sequence (GFP; negative control). Irrespective of the morpholino target, zygotes can hardly progress to the 8-cell stage at morpholino concentrations higher than 0.1 mM. Although this is suggestive of unspecific toxicity of morpholino no matter which target, it should be noted that targeting ZP2 is more detrimental than targeting GFP, as seen from the fact that the green bars in (A) are more extended to the right than the blue bars. At morpholino concentration of 0.05 mM both treatment and control zygotes progressed to blastocyst, however at rates substantially lower than 100% also in the negative control, suggestive of unspecific toxicity. At morpholino concentration of 0.01 mM majority of zygotes progressed to blastocyst in both treatment and control groups, indicating that this concentration is suited to test for gene-specific translation-blocking effects without incurring unspecific toxicity. Numbers of zygotes (n) microinjected with anti-ZP2 morpholino (mM concentration): 34 (0.01), 32 (0.05), 27 (0.1), 24 (0.5), 46 (1.0). Numbers of zygotes microinjected with control morpholino (mM concentration): 36 (0.01), 33 (0.05), 22 (0.1), 22 (0.5), 32 (1.0). (B). Zygotes microinjected with 0.01 mM morpholino were sampled at the 8-cell stage and subjected to ZP2 immunofluorescence using antibody ATCC cat.no. IE-3 – CRL-246. The immunofluorescence intensity values are provided in Supplementary Table S3. Immunofluorescence intensity quantification revealed that ZP2 intensity was reduced by ∼ 20% after translational inhibition of ZP2 mRNA compared to that of negative control. Representative images of the 8-cell embryos are shown after immunostaining with anti ZP2 (ATCC cat.no. IE-3 – CRL-246) revealed by secondary antibody. P-value was calculated with Wilcoxon test. *, <0.05. Abbreviations: IntDen, Integrated density of immunofluorescence signal (intensity x area).

### Protein KD reveals that intracellular ZP2 is required for the morula-to-blastocyst transition and healthy blastocyst formation

Given the ZP2 protein increase observed at the 8-cell / morula stage, we sought to determine its function by checking for effects when the increase is opposed. To this end, we used the Trim-Away method [12], and to account for translational ZP2 resynthesis (Figure 3), we combined Trim-Away with morpholino [13]. Trim-Away commits the protein of interest for proteasomal degradation using a specific antibody and the ubiquitin-protein ligase TRIM21. Morpholino prevents replenishment of the protein of interest by blocking its mRNA translation. To the best of our knowledge, this is the first time that the two methods have been applied together in mammalian embryos; we are only aware of a report in *Xenopus* where the two methods were applied separately [30]. In the following, unless otherwise stated, ‘ZP2 knockdown (ZP2 KD)’ refers to the combined treatment with Trim-Away and morpholino directed against ZP2; ‘mock knockdown (mock KD)’ refers to the combined treatment with Trim-Away and morpholino directed against an irrelevant protein (GFP in the case of our wildtype embryos); and ‘micromanipulation control (MC)’ refers to the treatment with Trim21 alone.

To induce ZP2 KD, zygotes were microinjected with a cocktail made of the Trim-Away reagents plus the *Zp2* morpholino, as per our established microinjection method [8, 28, 31]. The antibody was the same monoclonal antibody already used in immunofluorescence (ATCC cat.no. IE-3 – CRL-246), and the morpholino was the same as used before. The microinjected zygotes were cultured in KSOM(aa) medium and monitored daily to follow their developmental progression. ZP2 KD embryos looked retarded on day 3 and the morulae struggled to form blastocysts, compared to mock KD and MC groups (Figure 4A). At this stage, by proteomic analysis, the extent of ZP2 KD was estimated at 16% (Supplementary Figure S4), which is consistent with the previous results (Figure 3B). Closer examination of the blastocysts by triple immunofluorescence (CDX2, SOX17, NANOG) revealed that the ZP2 KD blastocysts were deficient in trophectoderm and primitive endoderm, compared to controls (Figure 4B). To understand if this was just a transient defect that could be compensated for at a later stage, ZP2 KD blastocysts were tested for postimplantation development. Compared to controls, ZP2 KD blastocysts had a significantly reduced ability to escape from the zona pellucida and attach to the feeder layer of fibroblasts in the outgrowth assay *in vitro*; *in vivo* they had a significantly reduced ability to develop to term after transfer to pseudopregnant uterus (Figure 4C).

**Figure 4.**
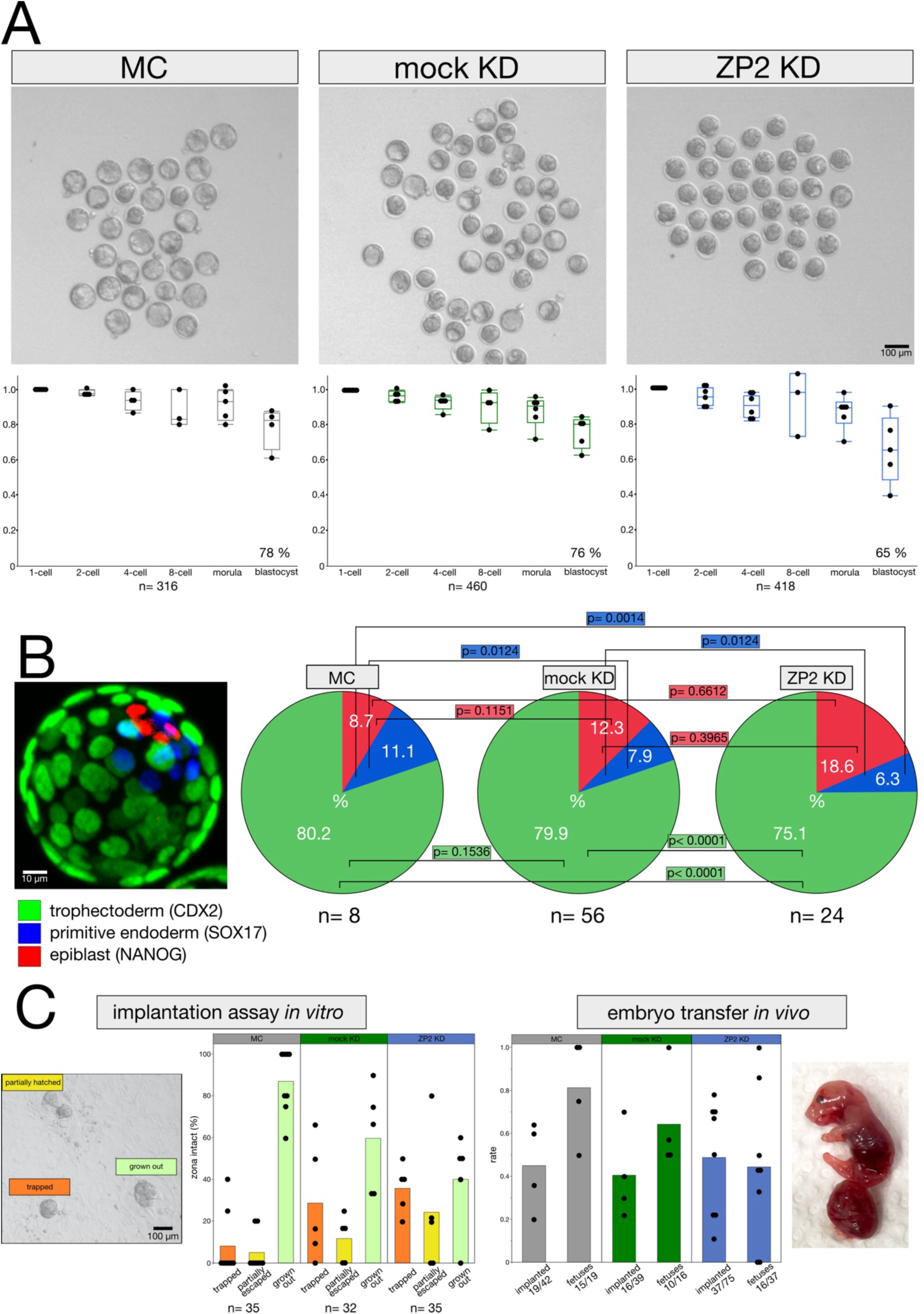
Intracellular ZP2 is required for the morula-to-blastocyst transition and healthy blastocyst formation. Zygotes were microinjected with protein KD reagents and cultured further. (A). At the morula stage (72h post fertilization), ZP2-KD embryos lagged behind in cavitation compared to mock KD and MC embryos. Numbers underneath the box plots are total number of embryos across the stages. The raw data of the developmental rates are provided in Supplementary Table S4. (B) At the blastocyst stage (96h post fertilization), ZP2-KD embryos were deficient in trophectoderm and primitive endoderm compared to mock KD and MC embryos. Numbers underneath the pie charts refer to the blastocysts imaged after triple immunofluorescence. The raw data of the cell counts are provided in Supplementary Table S5. (C) When examined for their ability to develop further, ZP2-KD blastocysts were less able of escaping the zona pellucida and forming outgrowths when plated onto a fibroblast layer *in vitro*; *in vivo* they were less able to develop to term when transferred to pseudopregnant uterus, compared to mock KD and MC blastocysts. Numbers underneath the bar plots are the blastocysts plated on feeders (implantation assay *in vitro*) or the fetuses obtained from the blastocysts transferred to uterus (embryo transfer *in vivo*). The raw data of the outgrowth and embryo transfers are provided in Supplementary Tables S6 and S7, respectively. P-values were calculated with Wilcoxon test. Abbreviations: ZP2 KD, ZP2 knockdown; mock KD, mock knockdown; MC, micromanipulation control.

The results of these KD experiments converge on an *interim* conclusion that intracellular ZP2 is not only present but also functionally active in mouse embryos. If ZP2 is depleted, then the morula-to-blastocyst transition is affected and the resultant blastocysts are unhealthy.

### Transcriptomic and proteomic correlates of unhealthy blastocyst formation after KD of ZP2

In search of a mechanistic explanation for the reduced ability of ZP2 KD embryos to form healthy blastocysts, we subjected these to transcriptome and proteome analysis. Let us briefly recall that the abbreviations ZP2 KD, mock KD and MC stand for ‘ZP2 knockdown’, ‘mock knockdown’ and ‘micromanipulation control’, respectively. MC consists of the injection of *Trim21* mRNA without the addition of antibodies or morpholino.

For transcriptome analysis we collected 10 blastocysts in triplicate (R1, R2, R3) for each of the treatments (ZP2 KD; mock KD; MC), and subjected them to RNA sequencing to measure the global gene expression levels (FPKM, fragments per kilobase of transcript per million mapped reads; see Methods). The dataset attained a depth of 37740 total RNAs (miRNA, snRNA, snoRNA, rRNA, lncRNA, protein-coding; Supplementary Tables S8), of which 17341 RNAs were present in all samples in all replicates (Supplementary Tables S9), and 12920 RNAs were protein-coding (Supplementary Tables S10). The full dataset has been deposited in ArrayExpress (E-MTAB-16339). The subset of 12920 mRNAs was used for further analysis. Non-hierarchical clustering showed that the samples clustered according to treatment (Figure 5A). Principal component analysis separated the ZP2 KD from the other two treatments in the 2nd component (Figure 5B). Volcano plots were applied to identify the mRNAs that are perturbed in the ZP2 KD or mock KD relative to MC (t-test, p<0.05; |fold change|>2; Figure 5C). Venn diagram intersections were applied on the two sets of perturbed mRNAs genes, to isolate the transcripts that are affected exclusively in the ZP KD or exclusively in the mock (Figure 5D). These mRNAs were subjected to overrepresentation analysis for ‘Biological Process’ by WebGestalt (see Methods). This analysis revealed that the top 10 terms in ZP2 KD are all significant (FDR <0.05) and include terms that are relevant to early post-implantation development (‘endoderm development’, ‘gastrulation’), while none of the top 10 terms are significant in case of mock KD (Figure 5D).

**Figure 5.**
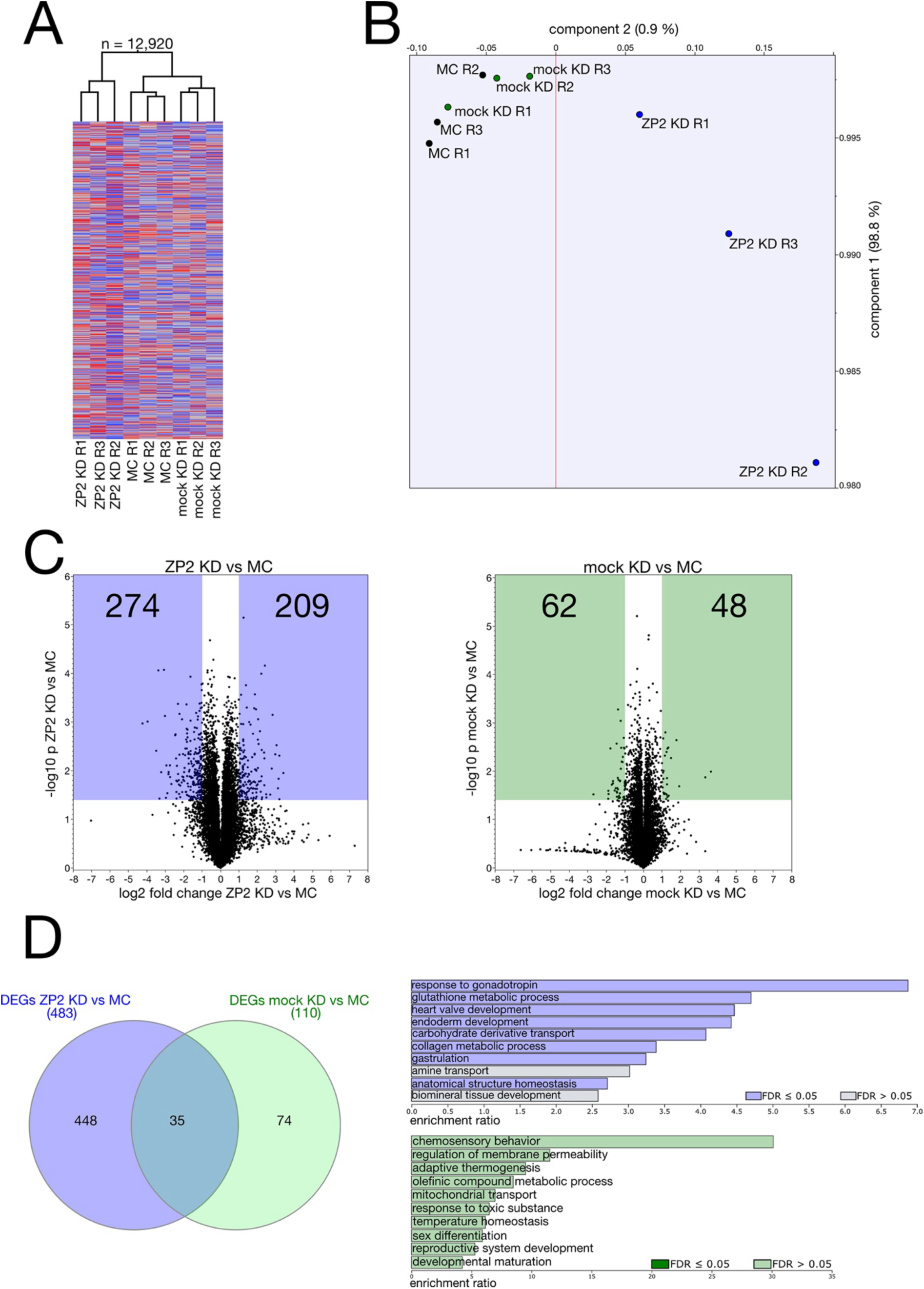
Transcriptomic correlates of unhealthy blastocyst formation after KD of ZP2. Zygotes were microinjected with protein KD reagents and cultured to blastocyst stage. (A) Hierarchical clustering of the 12920 protein-coding mRNAs detected in all samples. (B) Principal component analysis of the samples based on the protein-coding mRNAs. (C) Volcano plots of the mRNAs differentially expressed between ZP2 KD and MC and between mock KD and MC, highlighting the mRNAs that satisfy the criteria of both p<0.05 (Student’s t-test) and |fold change|ti2. The numbers in the colored areas indicate the respective differentially expressed mRNAs. (D) Venn diagram identification of mRNAs differentially exclusively in ZP2 KD and mock KD. Overrepresentation analysis of the mRNAs differentially expressed exclusively in ZP2 KD and mock KD in the ontology “Biological process”. Abbreviations: ZP2 KD, ZP2 knockdown; mock KD, mock knockdown; MC, micromanipulation control.

For proteome analysis we collected embryos at the late morula stage instead of blastocyst, so as to mitigate the risk of protein re-synthesis in case the morpholino had been consumed since the microinjection done three day before. We collected 30 late morulae in quadruplicate (R1, R2, R3, R4) for each of the treatments (ZP2 KD; mock KD; MC), and subjected them to mass spectrometric analysis in data-independent acquisition mode (see Methods) to gather the protein levels. The full dataset has been deposited in PRIDE (PXD062983). Overall, the dataset had a depth of 4531 proteins (Supplementary Table S11), of which 2401 were present in all samples in all replicates (Supplementary Table S12). The 2401 proteins dataset was used for further analysis. The samples clustered according to treatment (Figure 6A). Principal component analysis separated the ZP2 KD from the other two treatments by inspecting both the 1st and 2nd component (Figure 6B). Adopting the same workflow used for the transcripts, Volcano plots were applied to identify proteins that are perturbed in the ZP2 KD or mock KD relative to MC (t-test, p<0.05; |fold change|>2; Figure 6C); and Venn diagram intersections were applied to isolate the proteins that are affected exclusively in the ZP2 KD or exclusively in the mock KD (Figure 6D). These proteins were subjected to overrepresentation analysis in the ontology ‘Biological Process’ using WebGestalt. This analysis revealed that four of the top 10 terms in ZP2 KD are significant (FDR <0.05), and these four terms are related to protein synthesis, while none of the top 10 terms is significant in mock KD (Figure 6D).

**Figure 6.**
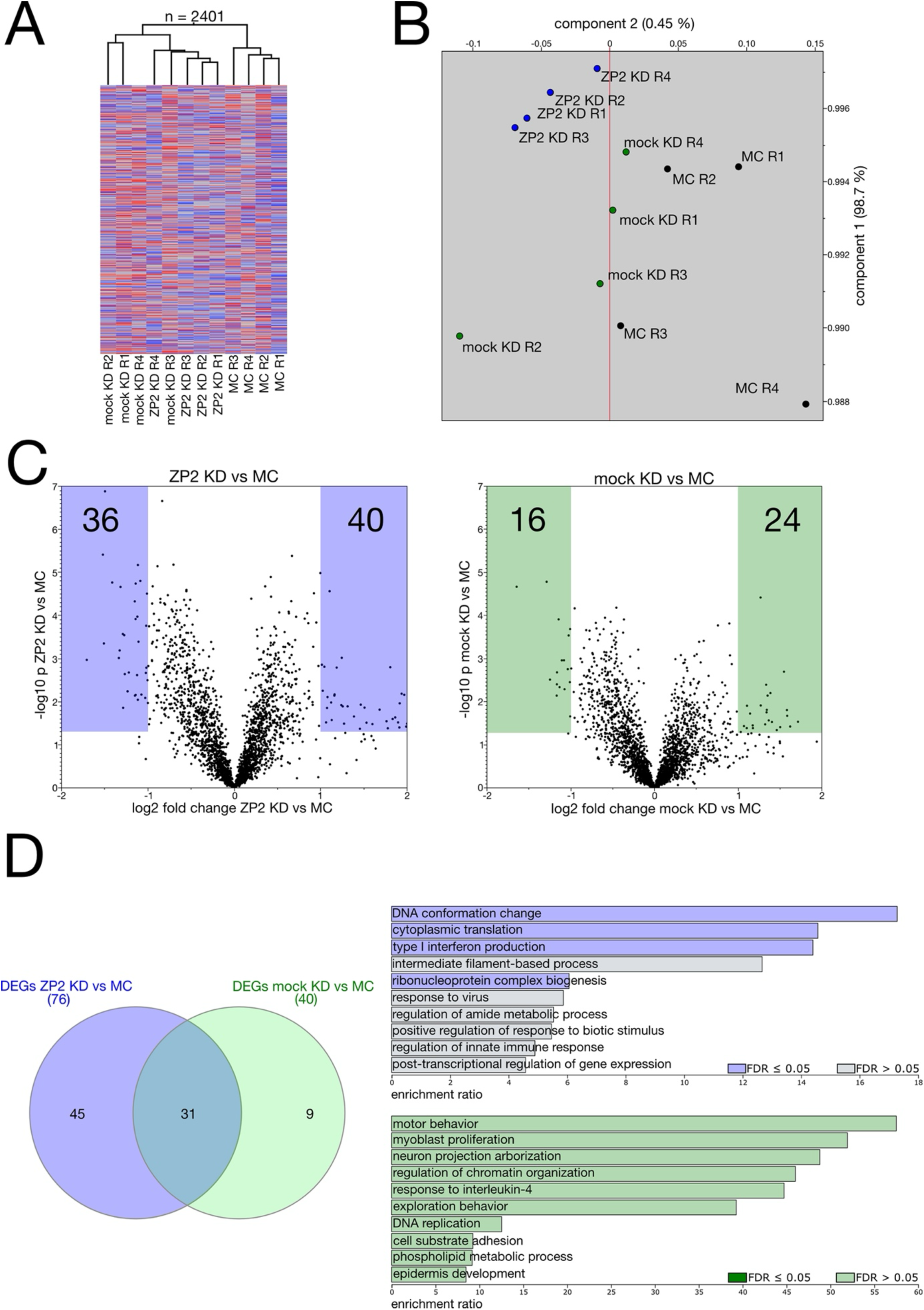
Proteomic correlates of unhealthy blastocyst formation after KD of ZP2. Zygotes were microinjected with protein KD reagents and cultured to later morula stage. (A) Hierarchical clustering of the 2401 proteins detected in all samples. (B) Principal component analysis of the samples based on the 2401 proteins. (C) Volcano plots of the proteins differentially expressed between ZP2 KD and MC and between mock KD and MC, highlighting the proteins that satisfy the criteria of both p<0.05 (Student’s t-test) and |fold change|ti2. The numbers in the colored areas indicate the respective differentially expressed proteins. (D) Venn diagram identification of proteins differentially exclusively in ZP2 KD and mock KD and mock. Overrepresentation analysis of the proteins differentially expressed exclusively in ZP2 KD and mock KD in the ontology “Biological process”. Abbreviations: ZP2 KD, ZP2 knockdown; mock KD, mock knockdown; MC, micromanipulation control.

The results of gene expression analysis are consistent with the phenotypic anomalies of ZP2 KD blastocysts described in Figure 4.

## Discussion

This study revealed that ZP2 protein, previously thought to solely be part of the secreted zona pellucida that surrounds oocytes and preimplantation embryos, is also present inside embryos, in mice. Furthermore, the intracellular amount of ZP2 increases at the 8-cell / morula stage and plays a functional role in blastocyst formation, specifically in trophectoderm and primitive endoderm formation. This novel and critical role may seem to go against a body of scientific literature about the known properties of the zona pellucida. In fact, it is not the first time that an increase in mouse embryos’ ZP protein content has been detected - the point is that no one had paid attention yet. For example, Gao and colleagues had also detected an increase, albeit not at the morula but at the blastocyst stage, using mass spectrometry (Table S1 in [32]). We demonstrated the increase using two orthogonal methods – mass spectrometry and immunofluorescence with monoclonal antibody. There was some variability in the stage when the increase was observed, which we attribute to the fact that naturally fertilized embryos are not perfectly synchronous, and the asynchrony is exacerbated by *in vitro* culture. Yet, what caused the ZP2 increase during embryogenesis, and is it spurious or does it have a function? As to the cause, the only possibility is that there is recruitment of residual mRNAs for protein synthesis, since *Zp2* transcripts do not increase during embryogenesis. As to the function, we reasoned that preventing translation should help illuminate the function, if it existed. The translation of mRNAs can be prevented using the morpholino method [13], which in our case opposed the increase of ZP2 observed at the 8-cell / morula stage. Since the newly synthesized quantity was added to a pre-existing quantity, we combined the morpholino method with the Trim-Away method, thereby targeting both quantities and producing a KD of ZP2. As a result we observed an impairment of the morula-blastocyst transition, which goes against the mainstream narrative about the role of ZP proteins. However, even in this case, it is not the first time that abnormal blastocyst formation has been observed, only that it was not discussed [5]. These findings will be discussed on three levels: 1) How to explain that we detected ZP proteins and particularly ZP2 inside mouse embryos when previous investigators did not? 2) what is the purpose of ZP2 protein re-synthesis in the embryo? 3) How to explain that the ZP2 KD embryos in this study suffered reduced post-implantation rates but survived, whereas embryos of *Zp2* KO oocytes died [5]? We provide the following answers to these questions as follows.

How to explain that we detected ZP proteins and particularly ZP2 inside mouse embryos when previous investigators did not? In principle it is not surprising that residual ZP proteins are still present in the embryo, assuming that secretion during oogenesis did not deplete the supply chain from the endoplasmic reticulum and Golgi, and considering that the half-life of the ZP proteins is long, being estimated as >100h [33, 34], which is approximately the duration of mouse pre-implantation development. But why did investigators before us not detect the intracellular ZP2? Since the zona is a secreted product that forms a coat around the cell, it always seemed nonsense to remove the very thing that one wanted to detect, and so the zona was left in place in previous studies. This way, however, the coat acted like a sponge that bound the antibody before this could penetrate deeper, ultimately obscuring the intracellular deposit of ZP proteins. A similar phenomenon has been described in a study showing the artefacts generated by indirect immunofluorescence when examining oocyte components located at the cell periphery, such as the subcortical maternal complex [35]. In our case, we made the same observation with both, direct and indirect immunofluorescence on specimens that were deprived of the zona, which is why our conclusion is safe: the ZP2 protein is internal, consistent with the results of mass spectrometry. As intriguing as the intracellular finding is the other finding about the subcellular localization. ZP2 should be located inside secretory vesicles, but if that were the case, then ZP2 should not be accessible to depletion by Trim-Away. Since the depletion occurred, it suggests that there is also ZP2 outside the vesicles, that is, in the cytosol. The presence of a cytosolic i.e. not intra-vesicular fraction was already demonstrated by us for ZP3 [8]. We speculate that the distinct subfractions may reflect distinct isoforms of ZP2. Such isoforms are already documented at the transcript and peptide levels of *Zp2* (Ensembl, Uniprot), while in terms of additional function and localization, the only known case is that of ZP3. Of note, Schultz and colleagues reported on a tumor cell-specific ZP3 transcript that encodes an intracellular cancer antigen [36].

What is the purpose of ZP2 protein re-synthesis in the embryo? We should like to acknowledge that the new function uncovered here had possibly already been ‘in the air’. In the case of the ZP3 partner of ZP2, Gerovska and Arauzo-Bravo reported that ZP3 has a burst of transcription in epiblast cells [37]. Epiblast is temporally and histologically not far away from the 8-cell / morula stage in which we detected ZP2 protein synthesis. There are several genes required for the morula to blastocyst transition in mice (*Sf3b14*, *Sf3b1, Rpl7l1*, and *Rrp7a* [38]; *Wdr74* [39]; *ZO-1* [40]; *Gm11545* [41]; *Tcfap2c* [42]; *Cdx2* [43]; *E-Cadherin* [44]; *EXOSC10/Rrp6* [45]; *Dcaf13* [46], *Traube (Trb)* a.k.a. *Aatf* [47]; *Actl6a*, *Gabpa*, *Hist1h3*, *Matr3*, *Mfng*, *Mxi1*, *Nop2*, *Pbrm1*, *Pnldc1*, *Ptpn18*, *Rpl7l1*, *Rrp7a*, *Rtn4*, *Sf3b1*, *Sf3b6*, *Supt6*, *Tm4sf1*, *Txnrd3*, *Uspl1*, and *Wdr74 [48]).* However, when the morula-stage requirement is additional to the canonical function, as in the case of *Zp2*, then there are not many genes known e.g. *Nr5a2*. This was previously known for its key role in mouse embryonic genome activation at the 2-cell stage [49] until it was also reported that it participates in morula stage viability [50]. The case of ZP2 is made even more interesting by the fact that not only is there a later expression stage (morula), but also a different subcellular localization, reminiscent of moonlight function. There are proteins that have a main function known to all (under the sunlight), but also a secondary function performed in another cell type or cell compartment that becomes visible when the sun is not shining (under the moonlight). There are many cases of moonlight proteins (reviewed in [51]), such as the signal transducer and activator of transcription 3 (STAT3), which has been found within mitochondria [52] in addition to its canonical role of regulating transcription in the nucleus to maintain the pluripotency of the blastocysts’ inner cell mass. It is noteworthy that these moonlight functions were often revealed via mass spectrometry in hypothesis-free studies that made it possible to notice that a protein was in the place where it was not expected, not to say the wrong place [53], just as happened to us when we first reported that there was zona pellucida within blastocysts [11].

How to explain that the ZP2 KD embryos in this study suffered reduced post-implantation rates but survived, whereas embryos of *Zp2* KO oocytes died [5]? Historically, the quesäon as to the biologic funcäon of gene products has been addressed by removing the DNA locus or its product e.g. mRNA to see what would happen without it. These approaches have already been applied to ZP2 [5, 18, 54], showing that the zona of growing oocytes does not increase in thickness if the *Zp2* mRNA is impaired during folliculogenesis [18, 54], and that the mutant oocytes of *Zp2* KO mothers do not support full development, with losses occurring in utero [5]. However, with protein KD, embryos deficient in ZP2 develop to term, albeit with difficulty at the blastocyst stage. We propose that the *Zp2* locus removed in the KO [5] precluded the increase of ZP2 at the 8-cell / morula stage, and that the paternal allele was insufficient compensate. Therefore, these embryos from KO oocytes had no option but to die, whereas in the case of the protein KD of this study there was compensatory ZP2 synthesis that made recovery – and embryo survival - possible.

To conclude, the message that emerges from this study is twofold. First, it documents that the silencing of oocyte-specific genes products at the transcriptional level can be overturned at the translational level. Second, it documents that a protein previously thought to mainly play a passive role outside the cell (physical protection) also plays an active role inside the cell (lineage formation). Pending cautious extrapolation to other species, our results expand the spectrum of considerations when advising subfertile women on the treatment of subfertility via medically assisted reproduction. When the zona is abnormal, bypassing the problem of polyspermic fertilization (thin zona pellucida) or fertilization failure (thick or hard zona pellucida) may well help the couple to produce embryos; but still these might be unable to produce normal blastocysts owing to the intracellular requirement of proteins such as ZP2 as uncovered here. Depending on which ZP protein is removed or mutated, this can lead to earlier or later blockade of embryonic cleavage, no matter if the zona defect was by-passed using ICSI. For these reasons it is tempting to recruit ZPs to the family of maternal factors [55] that are essential for early embryogenesis, at least in mice. At any rate, the case of ZP2 in this study documents that oocyte-specific genes can “speak up” again during embryogenesis, attesting to the biological complexity of embryogenesis that is likely to feature more surprising findings as technology develops further.

## Data availability

All data needed to evaluate this article are provided in the main body or in the Supplementary Materials. Large-scale data (omics) have been deposited in repositories. The mass spectrometry dataset of the developmental series presented in Figure 1 has been deposited in the PRIDE repository [56, 57] with accession number PXD056725 (“Application of SILAC method to the study of oocyte-to-embryo transition in mice”). For convenience this dataset is also available in processed form as Supplementary Table S1 on FigShare. The normalized integrated densities of ZP2 immunofluorescence intensities measured in pre-implantation embryos and presented in presented in Figure 2 are available in Supplementary Table S2 on FigShare. The data of the translational inhibition of Zp2 mRNA presented in Figure 3 are available in Supplementary Table S3 on FigShare. The rates of pre-implantation development, blastocyst quality and post-implantation development after ZP2 KD are presented in Supplementary Tables S4-S7. on FigShare. The RNA-seq dataset of Figure 5 has been deposited in the ArrayExpress with accession number E-MTAB-16339. For convenience this dataset is also available in processed form as Supplementary Tables S8-S10 on FigShare. The mass spectrometry dataset of Figure 6 has been deposited in the PRIDE repository with accession number PXD062983 (“Synthesis of oocyte-specific proteins during mouse embryogenesis and biological significance thereof”). For convenience this dataset is also available in processed form as Supplementary Tables S11-S12 on FigShare.

https://figshare.com/s/9108d71e4c1277e9372f

---

Supplementary Table S1_Fig 1_ Proteome of developmental series (riBAQ from MaxQuant after DDA)- PXD056725.xlsx

Supplementary Table S2_Fig 2_ Normalized integrated densities of ZP2 immunofluorescence on developmental series.xlsx

Supplementary Table S3_Fig 3_Morpholino test.xlsx

Supplementary Table S4_Fig 4A_Developmental rates after ZP2 KD, mock KD, MC.xlsx

Supplementary Table S5_Fig 4B_Cell lineage composition after ZP2 KD, mock KD, MC.xlsx

Supplementary Table S6_Fig 4C_Outgrowths after ZP2 KD, mock KD, MC.xlsx

Supplementary Table S7_Fig 4C_Embryotransfers after ZP2 KD, mock KD, MC.xlsx

Supplementary Table S8_Fig 5_37740 RNAs (all RNA categories)_ E-MTAB-16339.xlsx

Supplementary Table S9_Fig 5_17341 RNAs (all RNA categories present in all samples in all replicates, no zero values)_ E-MTAB-16339.xlsx

Supplementary Table S10_Fig 5_12920 mRNAs_no zero values_only protein coding_E-MTAB-16339.xlsx

Supplementary Table S11_Fig 6_4531 proteins (all values including zeroes) exp1350 PXD062983.xlsx

Supplementary Table S12_Fig 6_2401 proteins (non-zero values) exp1350 PXD062983.xlsx

## Acknowledgements

The authors express gratitude for scientific environment and infrastructural support to the Max Planck Institute for Molecular Biomedicine. As part of infrastructural support, we experienced outstanding support from the mouse housing facility, ensuring a dependable supply of mice needed to collect oocytes and embryos. The authors thank Annalen Buechsenschuetz and Saskia Winter for help with the LC-MS/MS measurements. The RNA-seq analysis was outsourced to the Core Genomic Facility of the Faculty of Medicine of the University of Muenster (Anika Witten, Andreas Huge). The authors also express gratitude to Irina Heinrich, who did her master degree in the Boiani group during the course of the experiments reported in this paper, contributing to some of them (immunoblots and IF imaging of embryos, embryo transfers).

## Authors’ roles

According to CRediT (Contributor Roles Taxonomy; https://credit.niso.org) M.B. and S.I. conceived the study (Conceptualization). M.B. acquired the funding (Funding acquisition). M.B. performed the micromanipulations, embryo culture and embryo transfers (Investigation) and analyzed the results (Formal Analysis). T.N. synthesized the mRNAs, purified the mRNAs and proteins for microinjection, did the immunoblots and the IF imaging, purified the mRNAs for RNA-seq (Investigation) and drew the figures (Visualization). H.D. performed the mass spectrometry analysis (Investigation). M.B. wrote the initial draft (Writing – original draft). M.B. and G.F. revised the subsequent versions of the manuscript (Writing – review & editing). All authors approved the manuscript and agree to be accountable for all aspects of the work.

## References

1. Zheng, P. and J. Dean. Oocyte-specific genes affect folliculogenesis, fertilization, and early development (2007). Semin Reprod Med. 25(4): p. 243–51.

2. Zeng, F. and R.M. Schultz. Gene expression in mouse oocytes and preimplantation embryos: use of suppression subtractive hybridization to identify oocyte- and embryo-specific genes (2003). Biol Reprod. 68(1): p. 31–9.

3. Epifano, O., et al. Coordinate expression of the three zona pellucida genes during mouse oogenesis (1995). Development. 121(7): p. 1947–56.

4. Roller, R.J., et al. Gene expression during mammalian oogenesis and early embryogenesis: quantification of three messenger RNAs abundant in fully grown mouse oocytes (1989). Development. 106(2): p. 251–61.

5. Rankin, T.L., et al. Defective zonae pellucidae in Zp2-null mice disrupt folliculogenesis, fertility and development (2001). Development. 128(7): p. 1119–26.

6. Lamas-Toranzo, I., et al. ZP4 confers structural properties to the zona pellucida essential for embryo development (2019). Elife. 8.

7. Fan, W., et al. Zona pellucida removal by acid Tyrode’s solution affects pre- and post-implantation development and gene expression in mouse embryosdagger (2022). Biol Reprod. 107(5): p. 1228–1241.

8. Israel, S., et al. Intracellular fraction of zona pellucida protein 3 is required for the oocyte-to-embryo transition in mice (2023). Mol Hum Reprod. 29(11).

9. Xiong, Z., et al. Ultrasensitive Ribo-seq reveals translational landscapes during mammalian oocyte-to-embryo transition and pre-implantation development (2022). Nat Cell Biol. 24(6): p. 968–980.

10. Zhang, H., et al. Stable maternal proteins underlie distinct transcriptome, translatome, and proteome reprogramming during mouse oocyte-to-embryo transition (2023). Genome Biol. 24(1): p. 166.

11. Taher, L., et al. The proteome, not the transcriptome, predicts that oocyte superovulation affects embryonic phenotypes in mice (2021). Sci Rep. 11(1): p. 23731.

12. Clid, D., et al. A Method for the Acute and Rapid Degradation of Endogenous Proteins (2017). Cell. 171(7): p. 1692–1706 e18.

13. Kanzler, B., et al. Morpholino oligonucleotide-triggered knockdown reveals a role for maternal E-cadherin during early mouse development (2003). Mech Dev. 120(12): p. 1423–32.

14. Percie du Sert, N., et al. The ARRIVE guidelines 2.0: Updated guidelines for reporting animal research (2020). PLoS Biol. 18(7): p. e3000410.

15. Summers, M.C., et al. IVF of mouse ova in a simplex optimized medium supplemented with amino acids (2000). Hum Reprod. 15(8): p. 1791–801.

16. Cox, J. and M. Mann. MaxQuant enables high peptide identification rates, individualized p.p.b.-range mass accuracies and proteome-wide protein quantification (2008). Nat Biotechnol. 26(12): p. 1367–72.

17. Schindelin, J., et al. Fiji: an open-source plajorm for biological-image analysis (2012). Nat Methods. 9(7): p. 676–82.

18. Tong, Z.B., L.M. Nelson, and J. Dean. Inhibition of zona pellucida gene expression by antisense oligonucleotides injected into mouse oocytes (1995). J Biol Chem. 270(2): p. 849–53.

19. Schwarzer, C., et al. ART culture conditions change the probability of mouse embryo gestation through defined cellular and molecular responses (2012). Hum Reprod. 27(9): p. 2627–40.

20. Boiani, M., et al. Oct4 distribution and level in mouse clones: consequences for pluripotency (2002). Genes Dev. 16(10): p. 1209–19.

21. Armant, D.R., H.A. Kaplan, and W.J. Lennarz. Fibronectin and laminin promote in vitro akachment and outgrowth of mouse blastocysts (1986). Dev Biol. 116(2): p. 519–23.

22. Elizarraras, J.M., et al. WebGestalt 2024: faster gene set analysis and new support for metabolomics and multi-omics (2024). Nucleic Acids Res. 52(W1): p. W415–W421.

23. Zhang, B., S. Kirov, and J. Snoddy. WebGestalt: an integrated system for exploring gene sets in various biological contexts (2005). Nucleic Acids Res. 33(Web Server issue): p. W741–8.

24. Liang, L.F., S.M. Chamow, and J. Dean. Oocyte-specific expression of mouse Zp-2: developmental regulation of the zona pellucida genes (1990). Mol Cell Biol. 10(4): p. 1507–15.

25. Bleil, J.D. and P.M. Wassarman. Structure and function of the zona pellucida: identification and characterization of the proteins of the mouse oocyte’s zona pellucida (1980). Dev Biol. 76(1): p. 185–202.

26. Shin, J.B., et al. Molecular architecture of the chick vestibular hair bundle (2013). Nat Neurosci. 16(3): p. 365–74.

27. Schwanhausser, B., et al. Global quantification of mammalian gene expression control (2011). Nature. 473(7347): p. 337–42.

28. Israel, S., et al. A framework for TRIM21-mediated protein depletion in early mouse embryos: recapitulation of Tead4 null phenotype over three days (2019). BMC Genomics. 20(1): p. 755.

29. Khosla, S., et al. Culture of preimplantation mouse embryos affects fetal development and the expression of imprinted genes (2001). Biol Reprod. 64(3): p. 918–26.

30. Chen, X., et al. Degradation of endogenous proteins and generation of a null-like phenotype in zebrafish using Trim-Away technology (2019). Genome Biol. 20(1): p. 19.

31. Israel, S., et al. The COP9 signalosome subunit 3 is necessary for early embryo survival by way of a stable protein deposit in mouse oocytes (2021). Mol Hum Reprod. 27(8).

32. Gao, Y., et al. Protein Expression Landscape of Mouse Embryos during Pre-implantation Development (2017). Cell Rep. 21(13): p. 3957–3969.

33. Shimizu, S., M. Tsuji, and J. Dean. In vitro biosynthesis of three sulfated glycoproteins of murine zonae pellucidae by oocytes grown in follicle culture (1983). J Biol Chem. 258(9): p. 5858–63.

34. Harasimov, K., et al. The maintenance of oocytes in the mammalian ovary involves extreme protein longevity (2024). Nat Cell Biol. 26(7): p. 1124–1138.

35. Jentod, I.M.A., et al. Mammalian oocytes store proteins for the early embryo on cytoplasmic lamces (2023). Cell. 186(24): p. 5308–5327 e25.

36. Schultz, I.J., et al. A tumor cell specific Zona Pellucida glycoprotein 3 RNA transcript encodes an intracellular cancer antigen (2023). Front Oncol. 13: p. 1233039.

37. Gerovska, D. and M.J. Arauzo-Bravo. Does mouse embryo primordial germ cell activation start before implantation as suggested by single-cell transcriptomics dynamics? (2016). Mol Hum Reprod. 22(3): p. 208–25.

38. Maserati, M., et al. Identification of four genes required for mammalian blastocyst formation (2014). Zygote. 22(3): p. 331–9.

39. Maserati, M., et al. Wdr74 is required for blastocyst formation in the mouse (2011). PLoS One. 6(7): p. e22516.

40. Wang, H., et al. Zonula occludens-1 (ZO-1) is involved in morula to blastocyst transformation in the mouse (2008). Dev Biol. 318(1): p. 112–25.

41. Kim, J., et al. Identification of a novel embryo-prevalent gene, Gm11545, involved in preimplantation embryogenesis in mice (2019). FASEB J. 33(10): p. 11326–11337.

42. Choi, I., et al. Transcription factor AP-2gamma is a core regulator of tight junction biogenesis and cavity formation during mouse early embryogenesis (2012). Development. 139(24): p. 4623–32.

43. Jedrusik, A., et al. Maternal-zygotic knockout reveals a critical role of Cdx2 in the morula to blastocyst transition (2015). Dev Biol. 398(2): p. 147–52.

44. Riethmacher, D., V. Brinkmann, and C. Birchmeier. A targeted mutation in the mouse E-cadherin gene results in defective preimplantation development (1995). Proc Natl Acad Sci U S A. 92(3): p. 855–9.

45. Petit, F.G., et al. EXOSC10/Rrp6 is essential for the eight-cell embryo/morula transition (2022). Dev Biol. 483: p. 58–65.

46. Zhang, Y.L., et al. DCAF13 promotes pluripotency by negatively regulating SUV39H1 stability during early embryonic development (2018). EMBO J. 37(18).

47. Thomas, T., et al. The murine gene, Traube, is essential for the growth of preimplantation embryos (2000). Dev Biol. 227(2): p. 324–42.

48. Cui, W., et al. Towards Functional Annotation of the Preimplantation Transcriptome: An RNAi Screen in Mammalian Embryos (2016). Sci Rep. 6: p. 37396.

49. Gassler, J., et al. Zygotic genome activation by the totipotency pioneer factor Nr5a2 (2022). Science. 378(6626): p. 1305–1315.

50. Festuccia, N., et al. Nr5a2 is dispensable for zygotic genome activation but essential for morula development (2024). Science. 386(6717): p. eadg7325.

51. Piatigorsky, J. Gene sharing in lens and cornea: facts and implications (1998). Prog ReIn Eye Res. 17(2): p. 145–74.

52. Tammineni, P., et al. The import of the transcription factor STAT3 into mitochondria depends on GRIM-19, a component of the electron transport chain (2013). J Biol Chem. 288(7): p. 4723–32.

53. Jeffery, C.J. Mass spectrometry and the search for moonlighting proteins (2005). Mass Spectrom Rev. 24(6): p. 772–82.

54. Pfender, S., et al. Live imaging RNAi screen reveals genes essential for meiosis in mammalian oocytes (2015). Nature. 524(7564): p. 239–242.

55. Innocenti, F., et al. Maternal effect factors that contribute to oocytes developmental competence: an update (2022). J Assist Reprod Genet. 39(4): p. 861–871.

56. Jones, P., et al. PRIDE: a public repository of protein and peptide identifications for the proteomics community (2006). Nucleic Acids Res. 34(Database issue): p. D659–63.

57. Perez-Riverol, Y., et al. The PRIDE database at 20 years: 2025 update (2025). Nucleic Acids Res. 53(D1): p. D543–D553.

