## Supplementary Figures and Tables for "A novel and critical role of the intracellular Zona Pellucida protein 2 (ZP2) for blastocyst formation in mice"

Supplementary Figure S1. Binding sites (A) and specificities (B) of ZP2 antibodies tested in this study.

Supplementary Figure S2. Immunofluorescent quantification of ZP2 in pre-implantation mouse embryos cultured in vitro.

Supplementary Figure S3. Quantification of ZP1, ZP2 and ZP3 transcript levels compared to those of housekeeping genes during mouse pre-implantation development.

Supplementary Figure S4. Quantification of ZP1, ZP2 and ZP3 proteins compared to housekeeping proteins may mass spectrometry at the morula/blastocyst stage (PRIDE PXD062983).

Supplementary Table S1_Fig 1_ Proteome of developmental series (riBAQ from MaxQuant after DDA)- PXD056725.xlsx

Supplementary Table S2_Fig 2_ Normalized integrated densities of ZP2 immunofluorescence on developmental series.xlsx

Supplementary Table S3_Fig 3_Morpholino test.xlsx

Supplementary Table S4_Fig 4A_Developmental rates after ZP2 KD, mock KD, MC.xlsx

Supplementary Table S5_Fig 4B_Cell lineage composition after ZP2 KD, mock KD, MC.xlsx

Supplementary Table S6_Fig 4C_Outgrowths after ZP2 KD, mock KD, MC.xlsx

Supplementary Table S7_Fig 4C_Embryotransfers after ZP2 KD, mock KD, MC.xlsx

Supplementary Table S8_Fig 5_37740 RNAs (all RNA categories)_ E-MTAB-16339.xlsx

Supplementary Table S9_Fig 5_17341 RNAs (all RNA categories present in all samples in all replicates, no zero values)_ E-MTAB-16339.xlsx

Supplementary Table S10_Fig 5_12920 mRNAs_no zero values_only protein coding_E-MTAB-16339.xlsx

Supplementary Table S11_Fig 6_4531 proteins (all values including zeroes) exp1350 PXD062983.xlsx

Supplementary Table S12_Fig 6_2401 proteins (non-zero values) exp1350 PXD062983.xlsx

### Supplementary Figures

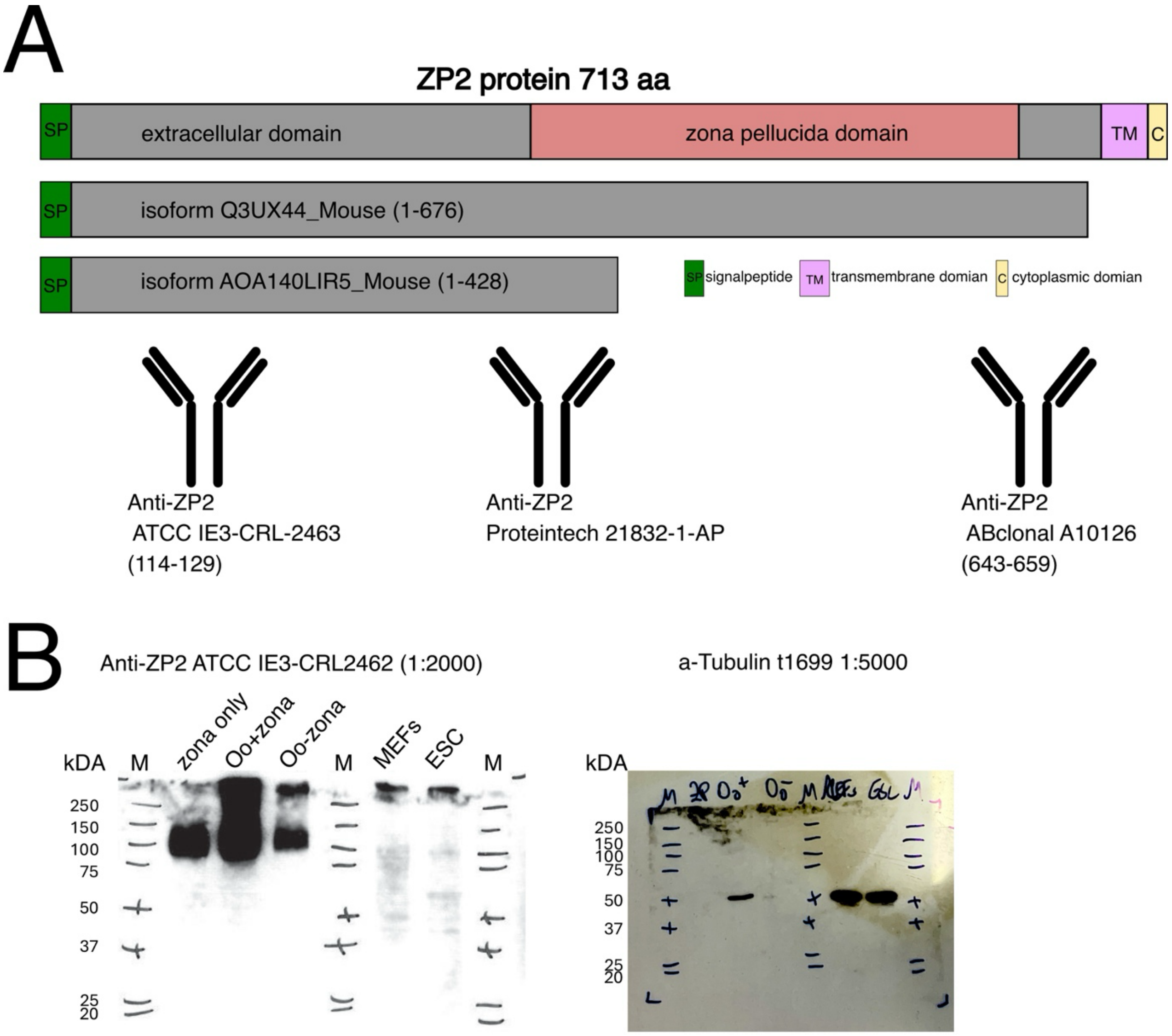


**Suppl. Figure S1. Antibody selection and validation.** Binding sites **(A)** and specificities **(B)** of ZP2 antibodies tested in this study. The same blot (B) was processed for ZP2, stripped and reprocessed for α-tubulin as the loading control.

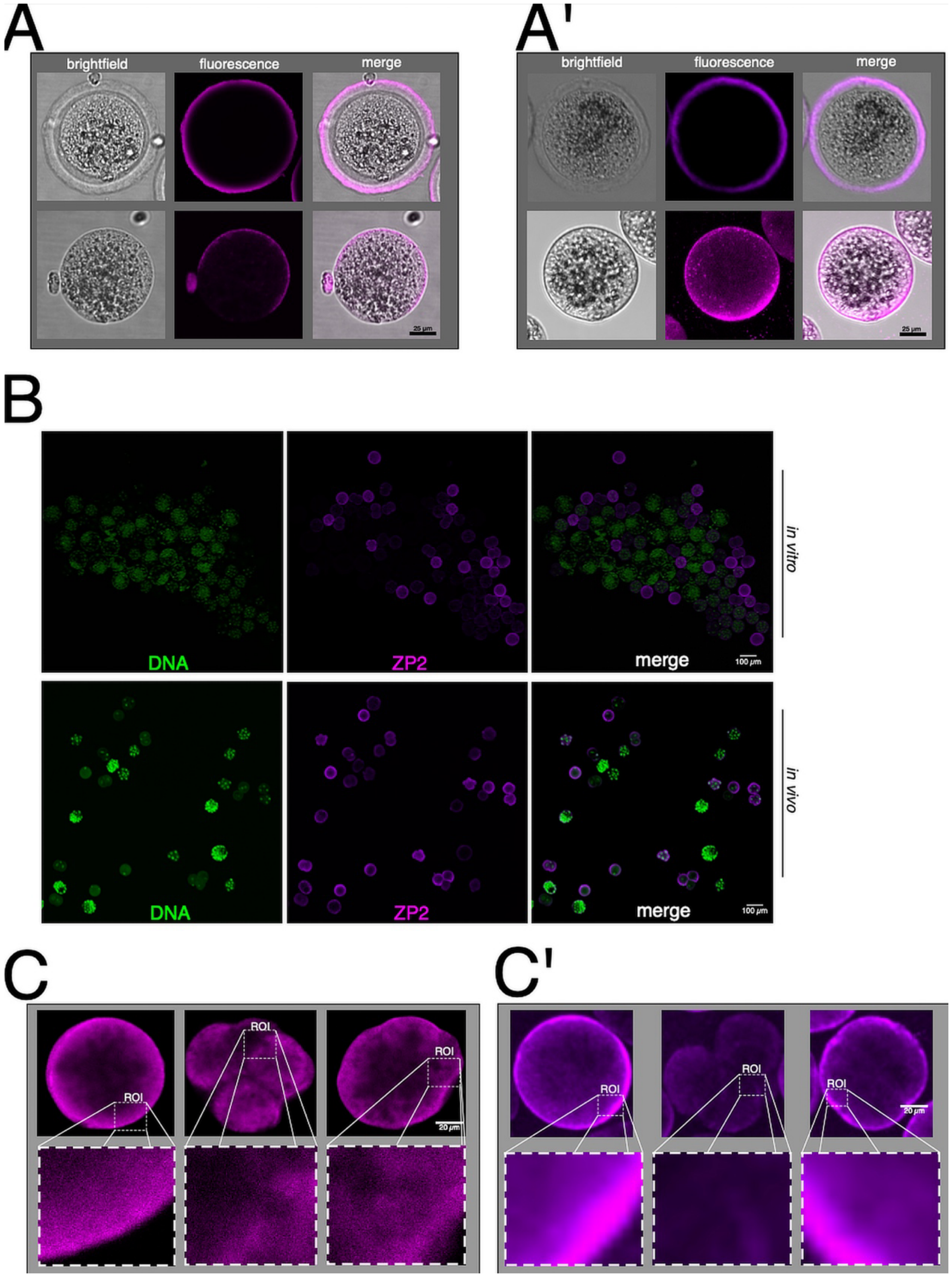


**Suppl. Figure S2. Immunofluorescence quantification of ZP2 in pre-implantation mouse embryos cultured in vitro. (A)**. Comparison of ZP2 signal intensity and distribution in zona intact and zona-free oocytes after indirect (A) and direct (A') immunofluorescence microscopy using ATCC cat.no. IE-3 – CRL-246 at the Dragonfly spinning disk microscope. **(B)**. Comparison of ZP2 signal intensity among preimplantation stages after indirect immunofluorescence with ATCC cat.no. IE-3 – CRL-246 at the Dragonfly spinning disk microscope. DNA was stained with YO-PRO-1. **(C)**. Super-resolution microscopy at the Zeiss LSM980 fitted with Airyscan 2 detector of zygotes (left), 4-cell embryos (center) and morulae (right) after indirect IF (C) and direct (C') immunofluorescence. The zona was removed post-fixation using acidic Tyrode's solution. Abbreviations: R.O.I., region of interest.

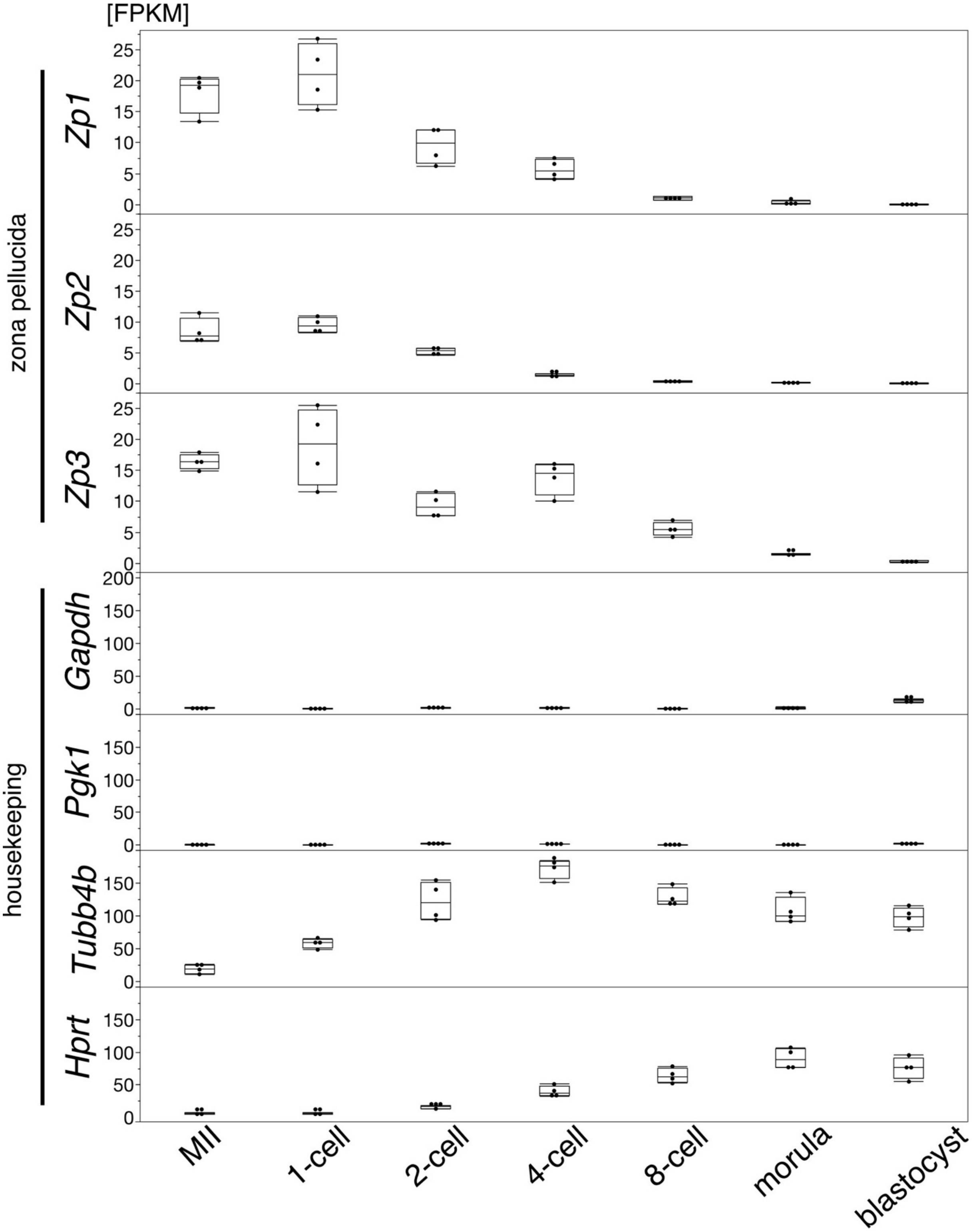


**Suppl. Figure S3. Quantification of ZP1, ZP2 and ZP3 transcript levels compared to housekeeping transcripts during mouse pre-implantation development.** Mouse embryos were collected at consecutive stages of pre-implantation development *in vitro* (culture in KSOM(aa) medium), deprived of the zona pellucida via Tyrode solution, and subjected to RNA-seq. Transcript levels are expressed as fragments per kilobase of transcript per million mapped reads (FPKM). The original dataset was published in Taher et al., (PMID 34887460).

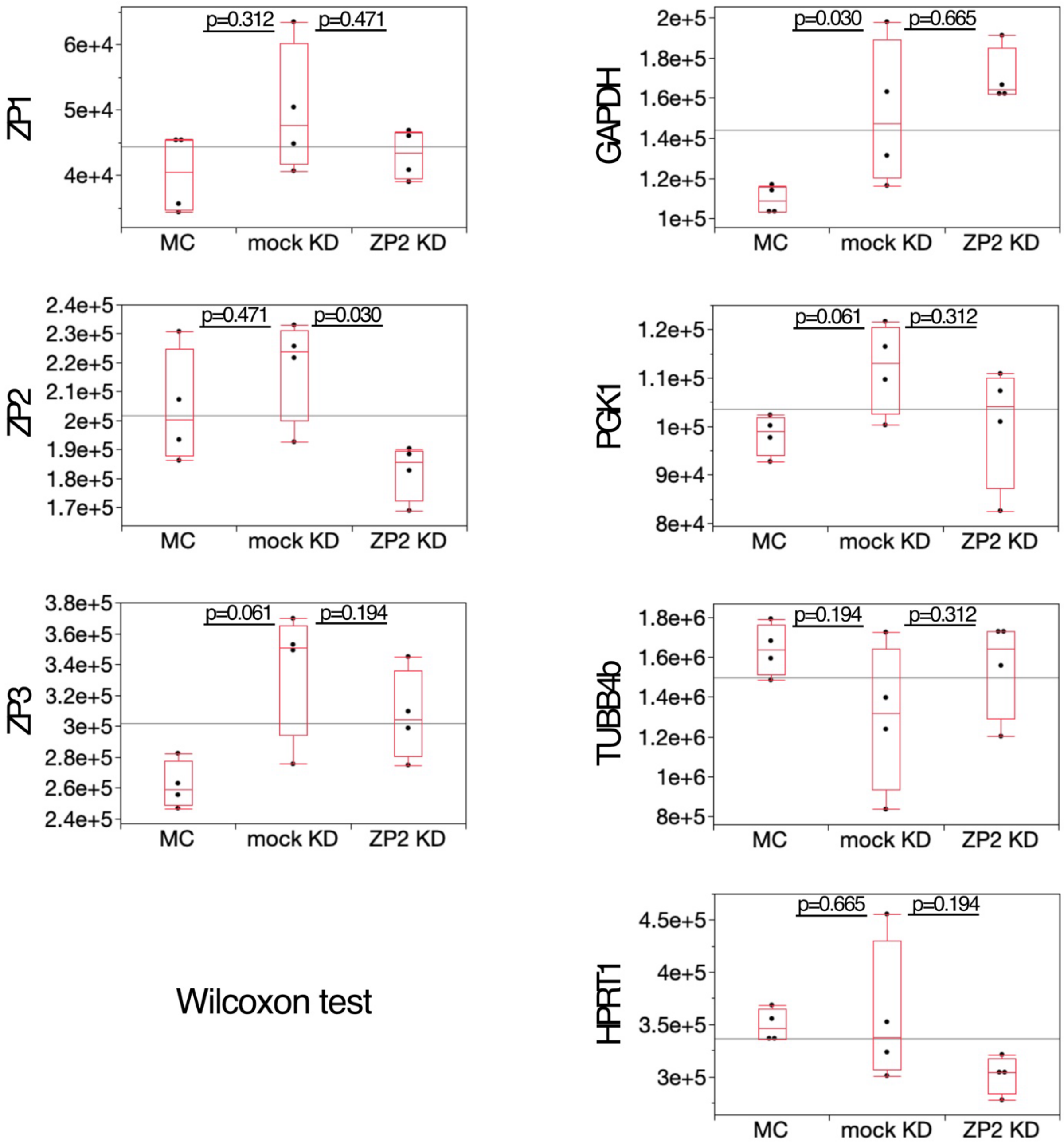


**Supplementary Figure S4.** Quantification of ZP1, ZP2 and ZP3 proteins compared to housekeeping proteins (GAPDH, PGK1, TUBB4b, HPRT1) by mass spectrometry at the late morula stage (dataset PRIDE PXD062983). Protein levels were quantified by DIA-NN. P-values were calculated with Wilcoxon test.

#### Supplementary Tables

Supplementary Table S1_Fig 1_ Proteome of developmental series (riBAQ from MaxQuant after DDA)- PXD056725.xlsx
Supplementary Table S2_Fig 2_ Normalized integrated densities of ZP2 immunofluorescence on developmental series.xlsx
Supplementary Table S3_Fig 3_Morpholino test.xlsx
Supplementary Table S4_Fig 4A_Developmental rates after ZP2 KD, mock KD, MC.xlsx
Supplementary Table S5_Fig 4B_Cell lineage composition after ZP2 KD, mock KD, MC.xlsx
Supplementary Table S6_Fig 4C_Outgrowths after ZP2 KD, mock KD, MC.xlsx
Supplementary Table S7_Fig 4C_Embryotransfers after ZP2 KD, mock KD, MC.xlsx
Supplementary Table S8_Fig 5_37740 RNAs (all RNA categories)_ E-MTAB-16339.xlsx
Supplementary Table S9_Fig 5_17341 RNAs (all RNA categories present in all samples in all replicates, no zero values)_ E-MTAB-16339.xlsx
Supplementary Table S10_Fig 5_12920 mRNAs_no zero values_only protein coding_E-MTAB-16339.xlsx
Supplementary Table S11_Fig 6_4531 proteins (all values including zeroes) exp1350 PXD062983.xlsx
Supplementary Table S12_Fig 6_2401 proteins (non-zero values) exp1350 PXD062983.xlsx

https://figshare.com/s/9108d71e4c1277e9372f
